# Extinction order affects community composition under strong competition and asymmetric species interactions

**DOI:** 10.64898/2026.10.22.761045

**Authors:** Joe Brennan, Sebastian J. Schreiber

**Author notes:** **Author Contributions Joe Brennan:** Conceptualization (lead), methodology (lead), software (lead), investigation (lead), writing - original draft, writing - review and editing (equal), visualization (lead). **Sebastian J. Schreiber:** Conceptualization (supporting), methodology (supporting), software (supporting), investigation (supporting), writing - review and editing (equal), visualization (supporting), supervision.

## Abstract

Extinction is a fundamental process impacting how community compositions change though time. Despite this, relatively few studies have examined how the order of species loss in multi-species extinction sequences can differentially impact the composition of the resulting disassembled community, known as an inverse priority effect. Here, we investigate the prevalence and predictors of inverse priority effects across a range of ecological contexts. First, we extend previous theory by deriving all possible cases of inverse priority effects in three-species Lotka-Volterra systems of coexisting competitors with equal intrinsic growth rates. We show that communities with strong, asymmetric competitive interactions are more likely to exhibit inverse priority effects. We then use a random matrix approach to demonstrate that inverse priority effects become more prevalent in more speciose communities and vary with the length of the extinction sequence. Furthermore, in speciose communities, inverse priority effects become increasingly likely with less facilitative or more competitive average interspecific interactions. Together, our results suggest that inverse priority effects are likely common in species-rich communities, particularly those with strong competition and asymmetric species interactions, warranting further theoretical and empirical investigation.

## Introduction

Extinction is a fundamental process that shapes the structure and functioning of ecological communities (Larsen et al., 2005). One community-level consequence of extinction is the loss of additional species following an initial extinction event, commonly termed a secondary extinction (Brodie et al., 2014). Secondary extinctions are suspected to contribute substantially to global biodiversity loss (Colwell et al., 2012; Strona and Bradshaw, 2018; Kehoe et al., 2021). Consequently, many studies have examined how species loss propagates through communities and which species are most likely to trigger secondary extinctions. Much of this work relies on static network models that do not explicitly incorporate population dynamics. In these frameworks, species are assumed to undergo secondary extinction when all or some proportion of their resources (e.g., prey or pollinators) are lost (Dunne et al., 2002, 2004). A common analysis in these studies examines how the order of species removal affects community disassembly, particularly the number of secondary extinctions that occur (Ávila-Thieme et al., 2023). These studies have provided insights into how the order of extinction (e.g., random versus empirically-predicted ordering) influences the magnitude of secondary extinction cascades (Berg et al., 2015).

While the order of species loss in static network models may generate different numbers of secondary extinctions, the final composition following disassembly is independent of the order of losses. This is because static network models account for secondary extinctions only when species lose required resources or mutualists (Memmott et al., 2004; Curtsdotter et al., 2011). Consequently, the final set of surviving species is determined by the set of species removed rather than their order of removal. Distinct disassembly outcomes require indirect effects mediated by population dynamics. Several empirical studies suggest that different sequences of species loss can produce distinct disassembly outcomes and levels of diversity in the resulting communities (Collins et al., 2009; Torres et al., 2024b). Thus, to holistically understand the impacts of extinction order on the outcomes of disassembly, explicit dynamical models of community dynamics are needed. Indeed, Torres et al. (2024a) illustrated that the order of extinction events can generate different disassembled communities in three-species competitive Lotka-Volterra models, a phenomenon termed an inverse priority effect. Moreover, they showed how niche and fitness differences can predict when inverse priority effects arise (Torres et al., 2024a).

A natural next step is to assess how common inverse priority effects are in more speciose Lotka-Volterra communities. However, the simulation-based technique for identifying inverse priority effects used in Torres et al. (2024a) would become highly computationally-intensive in speciose communities, especially if testing the impacts of different initial conditions at which extinction occurs. In contrast, recently developed theory uses invasion growth rates to identify transitions between community compositions due to species extinction (Brennan and Schreiber, 2026). For Lotka-Volterra systems, this approach only requires solving systems of linear equations to identify all possible transitions between community compositions due to extinction of a species. By studying the community disassembly graph that details all possible transitions between communities due to extinction, we can identify all possible inverse priority effects from a given community composition.

While the community disassembly graph can identify when inverse priority effects occur, it itself does not explain why inverse priority effects occur. To understand when inverse priority effects occur in three-species competitive communities, Torres et al. (2024a) used niche and fitness differences of the two-species subsets (Chesson and Kuang, 2008). However, niche and fitness differences do not describe higher order processes involving more than two species such as competitive asymmetry (Ranjan et al., 2024), a suspected mechanism for inverse priority effects (Torres et al., 2024a). Thus, we must consider alternate metrics describing ecological communities if we wish to extend the theory of inverse priority effects to more speciose communities.

Here, we extend the theory of inverse priority effects to more speciose communities. As a first step, we refine the analysis of Torres et al. (2024a) by classifying all possible inverse priority effects in three-species competitive Lotka-Volterra communities with equal intrinsic growth rates. We show that inverse priority effects are associated with strong, asymmetric competitive interactions. We then scale up our analysis to competitive communities with greater species richness and find that inverse priority effects become more likely in more speciose communities and with longer extinction sequences. Next, we examine communities containing both competitive and facilitative interactions and show that communities with relatively more competition than facilitation are more likely to exhibit inverse priority effects. Finally, using a machine learning approach, we demonstrate that variation in inverse priority effects is driven primarily by species richness and the number of species removed during the extinction sequence, emphasizing the probable presence of inverse priority effects in speciose, empirical systems.

## Methods

### Three species competitive communities

Torres et al. (2024a) studied the prevalence and predictors of inverse priority effects in the three-species competitive Lotka-Volterra model. In this model, species *i* grows exponentially at low density at rate *r*_*i*_ *>* 0 and species *j*’s per-capita competitive effect on species *i* occurs at rate *r*_*i*_*α*_*ij*_ *>* 0. Under these assumptions, the generalized Lotka-Volterra model is

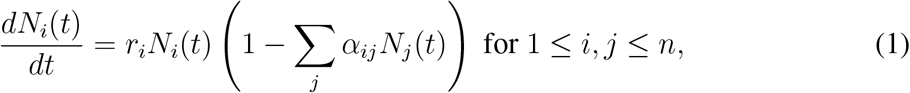

where, for three-species communities, *n* = 3. Through the change of variables *x*_*i*_ = *α*_*ii*_*N*_*i*_ (see Appendix A), Equation 1 simplifies to

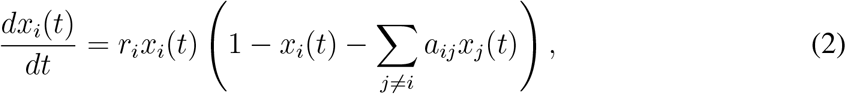

where 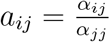 . As in Torres et al. (2024a), we assume all species have equal intrinsic growth rates (*r*_*i*_ = *r, ∀i ∈ {*1, …, *n}*). With this assumption, rescaling time such that *τ* = *rt* simplifies Equation 2 further to

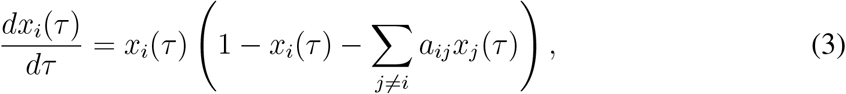

as derived in Appendix A.

With our model defined, we investigate how extinction order from a coexistence state influences the composition of the resulting disassembled community. There exist several choices for defining coexistence (Schreiber, 2006; Clark et al., 2026). Here, we define coexistence of the full community by permanence, characterized by a global attractor bounded away from extinction (Hofbauer and Sigmund, 1998; Schreiber, 2000). Because the three-species competitive Lotka-Volterra model is well-studied, all possible cases of three-species coexistence and their corresponding conditions for Equation 3 have been derived analytically (Hutson and Law, 1985; Zeeman, 1993; Jiang and Liang, 2020).

For permanent three-species competitive communities, there are four possible dynamical behaviors (Figure 1), with the corresponding parameter conditions provided in Appendix B. A feasible equilibrium of a subset of species is a strictly positive equilibrium of the system restricted to that subset (Clark et al., 2026). Across all classes, each species in isolation has such an equilibrium. Class 1 corresponds to the case in which every subset of species has a feasible equilibrium, corresponding to a biological system where competitive exclusion does not occur. Class 2 corresponds to the case where only two pairs of species have feasible equilibria. Biologically, class 2 corresponds to a system where one species is competitively excluded by another unless the third species is present, preventing competitive exclusion of the inferior species. Class 3 contains only one feasible two-species equilibrium, biologically representing a case where only one pair of species can coexist and, upon introduction of the third species, allows for coexistence of all three species. Class 4 has no feasible equilibria for any pair of species, representing strong competitive intransitivity wherein each species outcompetes another in a cyclic manner. Class 4 represents a coexistence scenario where mutual invasibility is not sufficient to guarantee three-species coexistence in the sense of permanence (May and Leonard, 1975; Schreiber, 2000; Chesson, 2018). For this class, we only use the parameter conditions that guarantee permanence (Jiang and Liang, 2020).

**Figure 1:**
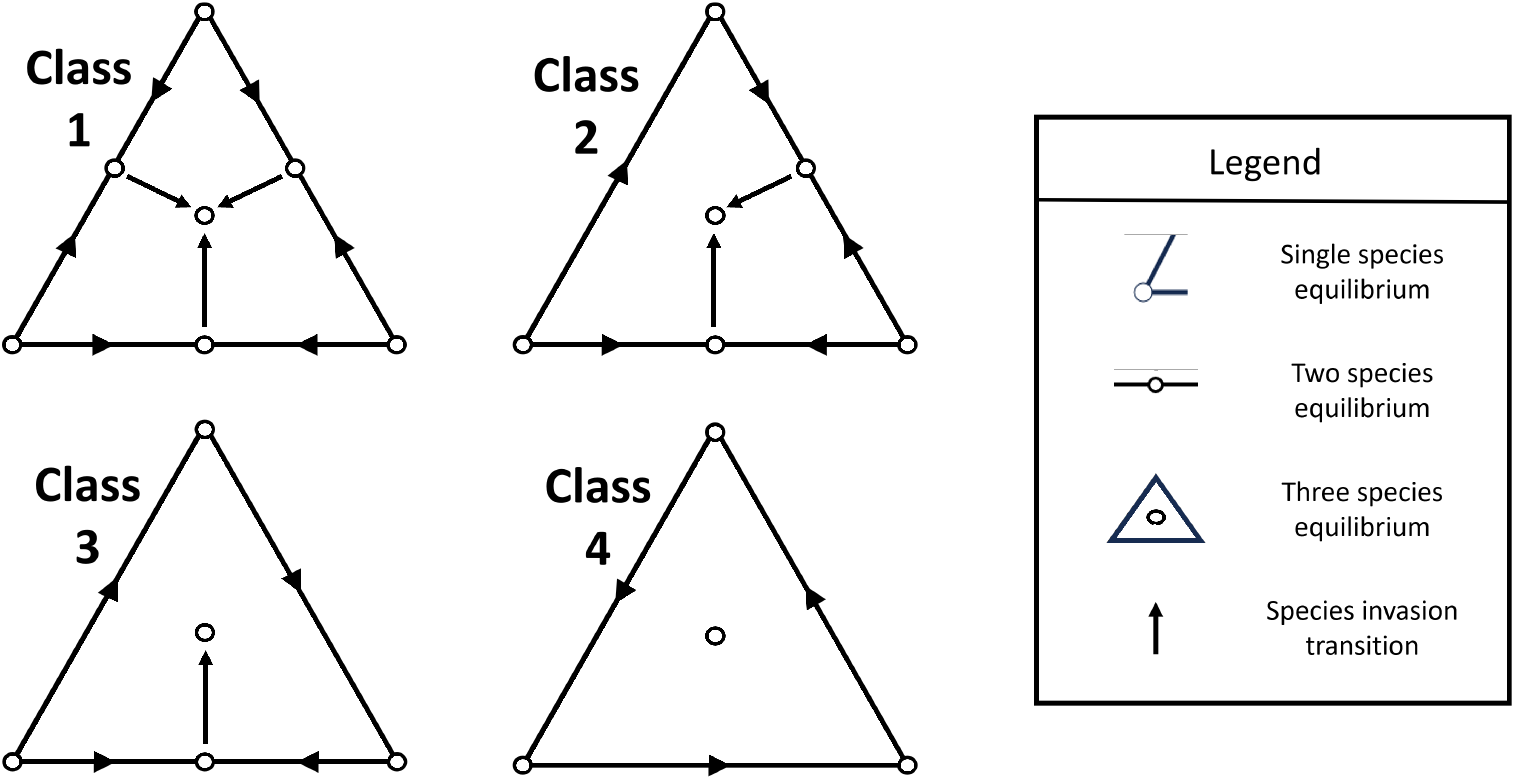
The four classes of dynamical behavior for the three-species competitive Lotka-Volterra model with equal intrinsic growth rates with three-species coexistence in the sense of permanence. Each triangle represents a phase portrait, displaying qualitative dynamics for each system. Vertices represent single species equilibria. Bottom left node denotes species 1, bottom right node denotes species 2, and top node denotes species 3. Edges represent when two species are at positive density, with a circle on the edge representing the existence of a feasible equilibrium for the pair of species. The interior of the triangle corresponds to all three species being present and the circle denotes the existence of a feasible equilibrium supporting all three species. A closed circle represents a stable equilibrium, open represents an unstable equilibrium, and half filled denotes a saddle point. Arrows denote transitions between equilibria due to introduction of a missing species.

Using this classification, we can systematically characterize all possible inverse priority effects in the system. To determine whether the order of removal of two species, say species *i* and *j*, generates an inverse priority effect, we compare the resulting disassembled communities under the two possible removal sequences. For a given dynamical class, we assume the system initially begins at the three-species coexistence state. We then remove one species (e.g., species *i*) and identify the resulting coexistence state. From this new state, we remove the second species (species *j*) and identify its new coexistence state. If species *j* is lost due to secondary extinction following the removal of species *i*, then the “removal” of species *j* in the second step has no impact on the composition of the community. We repeat this procedure for the reverse removal order. If the final disassembled communities differ between the two orderings, we classify the outcome as an inverse priority effect. We repeat this analysis for every pairwise combination of the three species in all four dynamical classes.

While this characterizes all possible cases of inverse priority effects in three-species competitive Lotka-Volterra systems, it does not identify the ecological mechanisms underlying them. In this setting, inverse priority effects require secondary extinctions; otherwise, the system converges to the single-species equilibrium of the species not removed. In competitive communities, asymmetric species interactions can increase the likelihood of secondary extinctions (Fowler, 2013; Brennan and Schreiber, 2026). Consequently, we hypothesize that three-species systems with greater asymmetry in competitive interactions are more likely to exhibit inverse priority effects (Torres et al., 2024a). To test this hypothesis, we generate 50,000 random three-species, competitive communities that coexist with interaction coefficients sampled such that *a*_*ij*_ = |*Z*_*ij*_|, *Z*_*ij*_ *∼ N* (0, 1). We use a half-normal distribution because its probability mass is concentrated near zero, reflecting the assumption that most species interact weakly with each other (Paine, 1992; O’Gorman et al., 2010). We then classify each interaction matrix according to the four dynamical classes in Figure 1.

For each interaction matrix, we calculate the average strength and asymmetry of species interactions. The average strength of species interactions equals 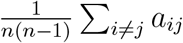 for a community of richness *n*. To define interaction asymmetry, let *A* be the interaction matrix whose (*i, j*) entry is *a*_*ij*_ in Equation 3. We decompose *A* into symmetric and antisymmetric components, 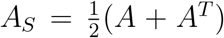 and 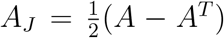, respectively. Interaction asymmetry is defined as 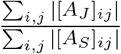 (Bortolan et al., 2025; *Calleja-Solanas et al*., *2026)*. A value of zero indicates nearly symmetric competitive interactions, while larger values correspond to increasingly asymmetric interactions. Evaluating interaction asymmetry across the different dynamical classes, we test whether communities exhibiting inverse priority effects have more asymmetric competitive interactions as hypothesized in previous studies (Torres et al., 2024a).

### More speciose competitive communities

To assess the prevalence of inverse priority effects in more speciose competitive communities, we use a random matrix approach. To do this, we randomly sample *a*_*ij*_ from a half normal distribution such that *a*_*ij*_ = |*Z*_*ij*_|, *Z*_*ij*_ *∼ N* (0, 1) to define the parameters of Equation 3. Our results are robust to other distributional assumptions (Appendix C). Once we sample all *a*_*ij*_, we test if the set of species can coexist in the sense of permanence using the invasion graph method (Hofbauer and Schreiber, 2022). Because we use the methodology described in Hofbauer and Schreiber (2022), we exclude parameterization with cycles in the invasion graph as there is no general characterization of permanence for higher dimensional systems with cycles (Hofbauer, 1994). However, our findings are qualitatively consistent if we instead allow a different form of coexistence that allows cycles but requires each disassembled community following removal of species *i* to be invadable by species *i*, reminiscent of classic tests of coexistence from Modern Coexistence Theory (Appendix C) (Chesson, 2000; Grainger et al., 2019). We allow subcommunities of the full, permanent community to coexist under a feasible equilibrium; though, requiring subcommunities to coexist under the stronger coexistence condition of permanence does not qualitatively change our findings (Appendix C). If all species coexist in the sense of permanence, we use the parameterization for our analyses. If not, we discard the parameterization and resample. We repeat this procedure 500 times for communities containing between three and six species, yielding a total of 2,000 parameterizations for comparison.

To identify inverse priority effects in these communities, we use the community disassembly graph (Brennan and Schreiber, 2026). Here, vertices of the disassembly graph represent feasible equilibria and directed edges with label *i* represent transitions from one community to another due to the extinction of species *i*. For Lotka-Volterra models, Brennan and Schreiber (2026) showed that the community disassembly graph can be obtained by solving for the invasion scheme (Hofbauer and Schreiber, 2022): all feasible equilibria of the system and the associated per-capita growth rates of all species at these feasible equilibria. For Equation 3, the percapita growth rate of species *i* at a feasible equilibrium supporting the species *S ⊂ {*1, …, *n}* is

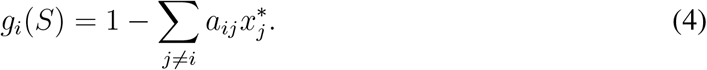

For the extinction of species *i* to cause a transition from community *S* to community *T*, two conditions must hold. First, the resulting community must satisfy *T ⊆ S \ {i}* meaning that *T* cannot contain species *i* and must be a subset of the original community after the removal of species *i*. Second, every species *j ∈ S \* (*T ∪ {i}*) must satisfy *g*_*j*_(*T* ) *<* 0. In other words, any species present in *S* but absent from *T*, aside from the initially removed species *i*, must have a negative invasion growth rate at community *T*. Species *j ∈ S \ {i}* who have a negative invasion growth rate at community *T* are species that experience secondary extinction following the loss of species *i* (Brennan and Schreiber, 2026). Repeating this procedure for all species *i ∈ S* and across all feasible communities defines the directed edges of the community disassembly graph.

Because the community disassembly graph details all possible transitions between community compositions due to extinction, it can identify all possible inverse priority effects. To determine whether an inverse priority effect arises for a given extinction order of the community *S*, we track the corresponding disassembly trajectory under each possible ordering of species loss, as illustrated in Figure 2. Specifically, given a set of species *R ⊂ {*1, …, *n}* in some extinction sequence, we consider every permutation *σ* : *R → R*. We then identify the disassembly path(s) associated with each permutation and record the set of final community composition(s) *S*(*σ*). An inverse priority effect occurs if *S*(*σ*_1_)≠ *S*(*σ*_2_) for at least one pair of permutations *σ*_1_*≠ σ*_2_.

**Figure 2:**
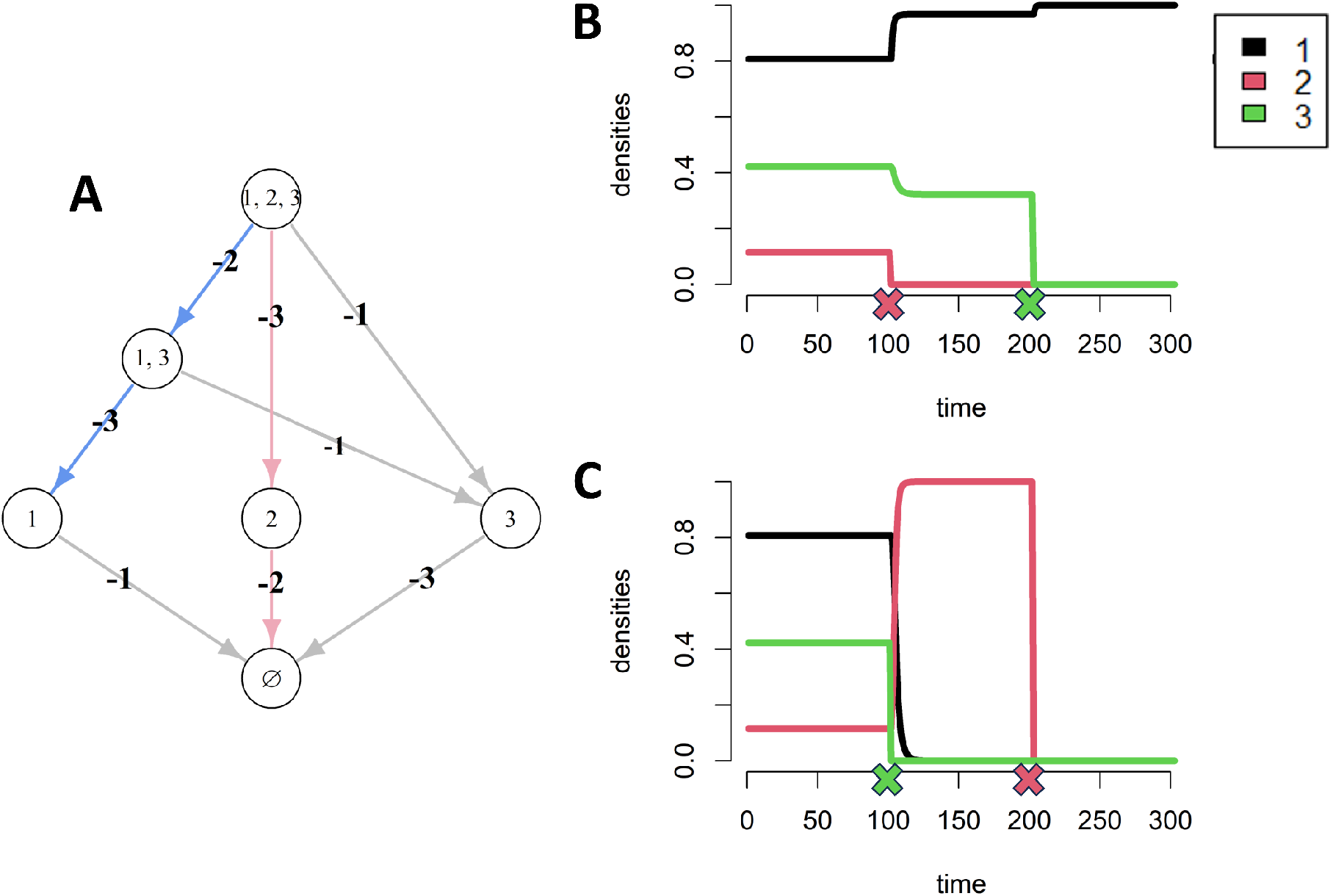
Using the community disassembly graph to identify inverse priority effects. (A) An example of the community disassembly graph calculated using invasion growth rates. The order in which species 2 and 3 are removed lead to different disassembled communities, corresponding to an inverse priority effect. (B) Simulation of the community dynamics with removal of species 2 followed by species 3. Colored “x” in the simulation denotes removal of the respective species at the corresponding timepoint. (C) Simulation of the community dynamics with removal of species 3 followed by species 2. Colored “x” in the simulation denotes removal of the respective species at the corresponding timepoint. Parameters are the same as the simulations in Torres et al. (2024a) where *a*_12_ = 1.3, *a*_21_ = *a*_32_ = *a*_13_ = 0.1, *a*_31_ = 0.7, and *a*_23_ = 1.9 for Equation 3.

For each simulated parameterization, we identify all possible inverse priority effects. Because we now consider communities containing more than three species, we also examine extinction sequences involving more than two species. Specifically, for a community of size *n*, we consider extinction sequences with lengths ranging from 2 to *n −* 1 species. In species rich systems, inverse priority effects can arise through two means: (i) at least one extinction sequence contains secondary extinctions or (ii) the removal of a species in an extinction sequence can lead to different resulting communities, corresponding to multiple directed edges in the community disassembly with same label *i* and the same starting community *S*. Once all possible inverse priority effects have been identified across parameterizations and richness levels, we evaluate how the probability of an inverse priority effect for a given extinction sequence depends on four metrics: species richness, proportion of species in the extinction sequence, interaction asymmetry, and average strength of interspecific competition. Additionally, we test an interactive effect between the average strength of interspecific competition and interaction asymmetry, as we hypothesize that communities with lower average competition will require more asymmetry for inverse priority effects to occur than more competitive communities.

To disentangle the effects of these variables on the probability of observing an inverse priority effect, we fit a binomial generalized additive model using the *mgcv* package in R (Version 4.5.0) (Wood and Wood, 2015). Because our simulation study is large, formal significance tests are not informative, since even negligible effects can be statistically significant. We therefore focus on the magnitude and direction of estimated effect sizes.

### Speciose communities with competition and facilitation

Next, we examine how facilitation influences the prevalence of inverse priority effects. As before, we randomly draw parameters for Equation 3, but now allow both negative (facilitative) and positive (competitive) values of *a*_*ij*_. Specifically, we sample *a*_*ij*_ *∼ N* (0, 1) to parameterize Equation 3. Although we present results only for normally distributed parameters, our conclusions are robust to alternative distributional assumptions (Appendix D). Since facilitation in Lotka-Volterra systems can lead to unbounded growth (Goh, 1979), after sampling interaction coefficients, we verify boundedness of all species combinations using the B-matrix condition described in Hofbauer and Sigmund (1998) using the algorithm presented in Schreiber et al. (2026). If the B-matrix condition is not satisfied, we discard the parameterization; otherwise, we then use the invasion graph approach to assess if the community containing all species coexists under permanence (Hofbauer and Schreiber, 2022). To ensure permanence, we do not consider parameterizations that generate cyclic invasion graphs, as one cannot easily verify permanence in such cases (Hofbauer, 1994). However, our findings are qualitatively consistent if we instead allow a different form of coexistence that permits cycles in the invasion graph by simply checking if each species *i* can invade the disassembled community following its removal (Appendix D). Additionally, we again allow subcommunities of the full, permanent community to coexist under a feasible equilibrium; though, requiring subcommunities to coexist in the sense of permanence does not qualitatively change our findings (Appendix D). We use the random parameterization if it meets all of these criteria. Since random parameterizations with facilitation are more likely to meet our criteria, we expand our species richness range to three to seven for this set of analyses and again collect 500 parameterizations for each richness level. To identify inverse priority effects, we again use the community disassembly graph and fit generalized additive models to disentangle the effects of four predictors of interest: species richness of the original community, the proportion of species included in the extinction sequence, interaction asymmetry, and the mean interspecific interaction strength. Again, we test an interactive effect between the average strength of interspecific interactions and interaction asymmetry, as we hypothesize that communities whose average interaction is facilitative will require more asymmetry for inverse priority effects to occur than those whose average interaction is competitive.

In addition to the generalized additive model, we employ a random forest algorithm using the *randomForest* package in R to assess the relative importance of predictors in explaining variation in the probability of inverse priority effects (Liaw and Wiener, 2002). Predictors with higher importance values contribute more strongly to variation in inverse priority effects. We include five predictors: species richness of the original community, the proportion of species in the extinction sequence, interaction asymmetry, mean interspecific interaction strength, and a binary predictor distinguishing communities that are purely competitive from those that include both competitive and facilitative interactions. To test importance, we evaluate two metrics of model performance: mean decrease in accuracy and mean decrease in Gini. Mean decrease in accuracy measures the reduction in out-of-bag prediction accuracy when the values of a predictor are randomly permuted, whereas mean decrease in Gini measures the total reduction in Gini impurity resulting from splits on that predictor across all classification trees (Liaw and Wiener, 2002). We then supplement this analysis with recursive partitioning trees, classifying the occurrence of inverse priority effects based on predictor thresholds and their interactions (Therneau et al., 2015).

## Results

### Three species competitive communities

All three-species competitive communities have at least one extinction sequence that has an inverse priority effect except for Class 1 (see Figure 1 for classes; Appendix E). As all species pairs coexist in Class 1, inverse priority effects are not possible. We find that systems with at least one inverse priority effect (Classes 2,3,4) have more asymmetric and stronger competitive interactions than those with none (Class 1) (Figure 3). Moreover, classes of behavior with more extinction sequences corresponding to inverse priority effects have stronger interaction asymmetry and competitive interactions than classes with less inverse priority effects (Appendix E).

**Figure 3:**
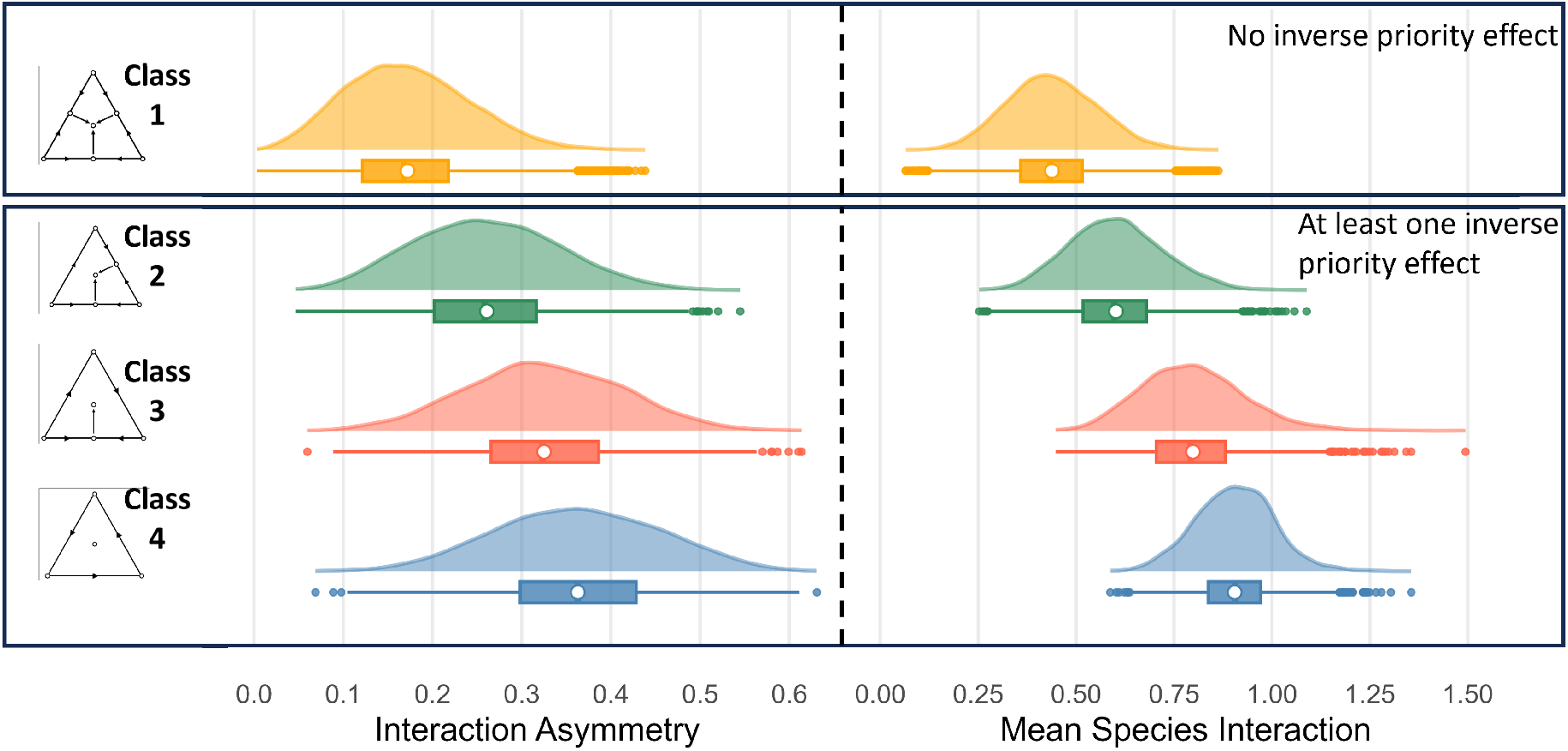
Inverse priority effects occur in communities with strong, asymmetric interactions. Density plots of asymmetry of simulated parameterizations with no inverse priority effects and at least one inverse priority effect (one, two, and three inverse priority effects for class 2, 3, and 4, respectively). Each row denotes a different dynamic classes 1 *−* 4, from top to bottom respectively. The left column shows each class’s interaction asymmetry values while the right column shows the mean species interaction strength. Below each density plot is a boxplot with the white circle denoting the mean value.

### More speciose, competitive communities

Asymmetric and strongly interacting communities are more likely to have an inverse priority effect (Figure 4A,B), qualitatively matching trends from the three species community (Figure 3). The probability of an inverse priority effect increases rapidly with interaction asymmetry at low asymmetry values. However, beyond a threshold, further increases in asymmetry do not substantially change this probability (Figure 4A). Similarly, as the mean strength of interspecific competition increases, the probability of an inverse priority effect also increases. However, in this case, the probability does not saturate with higher mean interaction strengths (Figure 4B). As hypothesized, communities with relatively low average interaction strength require more asymmetry for inverse priority effects to arise than communities with relatively high average interaction strength (Figure 4C). The probability of an inverse priority effect increases approximately linearly with species richness for the range of richnesses explored (Figure 4D). As the proportion of species removed (our normalized proxy for the length of the extinction sequence) increases, the probability of an inverse priority effect also increases, eventually saturating at higher values (Figure 4E, Appendix G).

**Figure 4:**
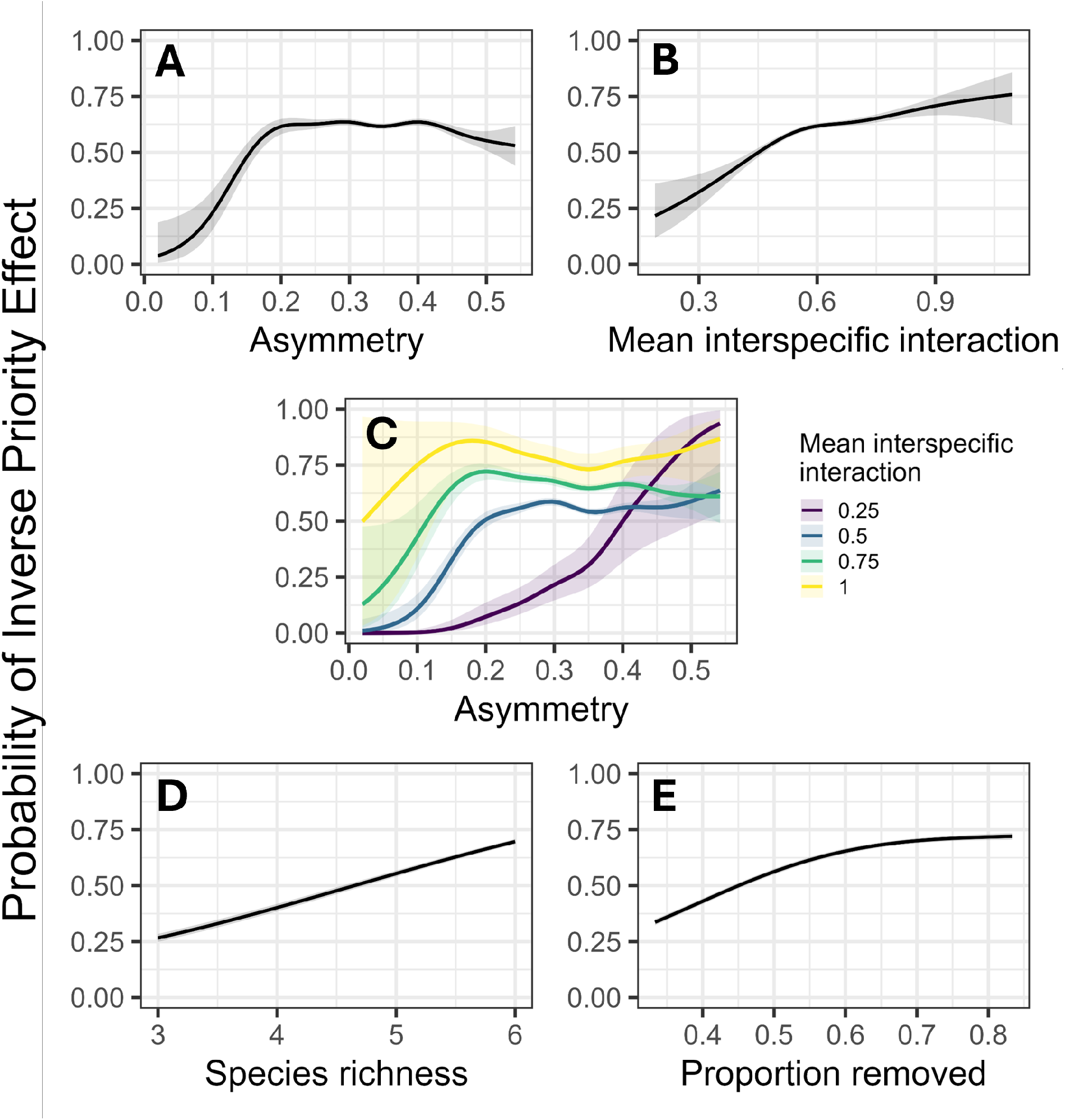
Predictors of inverse priority effects in competitive communities. (A) The conditional predicted probabilities of an inverse priority effect across the range of observed asymmetry values, with all other predictors held at their observed means. (B) The conditional predicted probabilities of an inverse priority effect across the range of observed mean interspecific interaction values. (C) The conditional predicted probabilities of an inverse priority effect when both asymmetry and mean interspecific interaction strength vary. (D) The conditional predicted probabilities of an inverse priority effect across the range of observed species richness values. (E) The conditional predicted probabilities of an inverse priority effect across the range of observed proportion of species removed values.

### Speciose communities with competition and facilitation

In communities with both competition and facilitation, small increases in asymmetry at relatively low values drive large increases in the probability of an inverse priority effect, while, at higher values, small increases negligibly change the probability of an inverse priority effect (Figure 5A). As the average interaction strength in the community becomes more competitive and less facilitative (i.e., as *a*_*ij*_ becomes more positive in Equation 3), the probability of an inverse priority effect increases (Figure 5B). Additionally, communities whose species are facilitative on average require more asymmetry to experience inverse priority effects than communities whose average interaction is competitive (Figure 5C). As species richness increases, we again observe an approximately linear increase in the probability of an inverse priority effect (Figure 5D). Increasing the proportion of species removed in the extinction sequence leads to an initial increase in the probability of an inverse priority effect, followed by a slight decline at higher values (Figure 5E). These relationships are qualitatively similar to the model with only competition (Figure 4).

**Figure 5:**
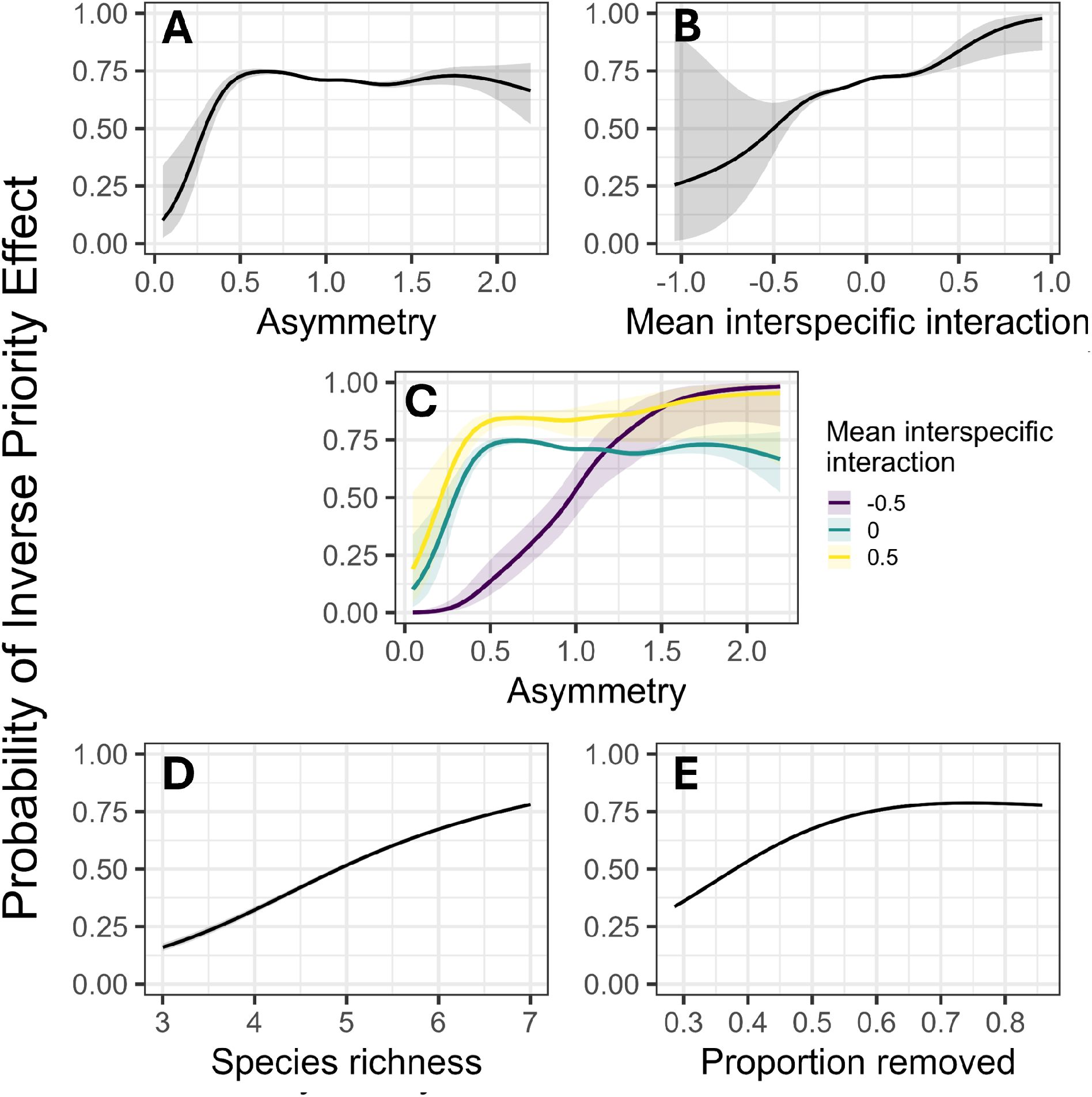
Predictors of inverse priority effects in communities with competition and facilitation. (A) The conditional predicted probabilities of an inverse priority effect across the range of observed asymmetry values, with all other predictors held at their observed means. (B) The conditional predicted probabilities of an inverse priority effect across the range of observed mean interspecific interaction values. Positive values correspond to competition while negative values correspond to facilitation. (C) The conditional predicted probabilities of an inverse priority effect when both asymmetry and mean interspecific interaction strength vary. (D) The conditional predicted probabilities of an inverse priority effect across the range of observed species richness values. (E) The conditional predicted probabilities of an inverse priority effect across the range of observed proportion of species removed values.

The importance values from the random forest analysis indicate that variation in species richness and variation in the proportion of the community included in the extinction sequence account for the most variability in inverse priority effect outcomes (Figure 6A,B). In contrast, whether the community contains only competitive interactions or a mixture of competitive and facilitative interactions does not substantially influence the variability of inverse priority effects. Instead, the interaction structure influenced by the asymmetry of the interactions and the mean interaction strength plays a more prominent role. In particular, further analyzes using recursive partitioning trees suggest that a stronger and more competitive mean interspecific interaction is an important predictor in lower richness communities (Figure 6C). These results further support the findings of the generalized additive models, suggesting that the qualitative effects of predictors on the probability of inverse priority effects are broadly consistent between purely competitive communities and those that include both competition and facilitation (Figures 4 and 5).

**Figure 6:**
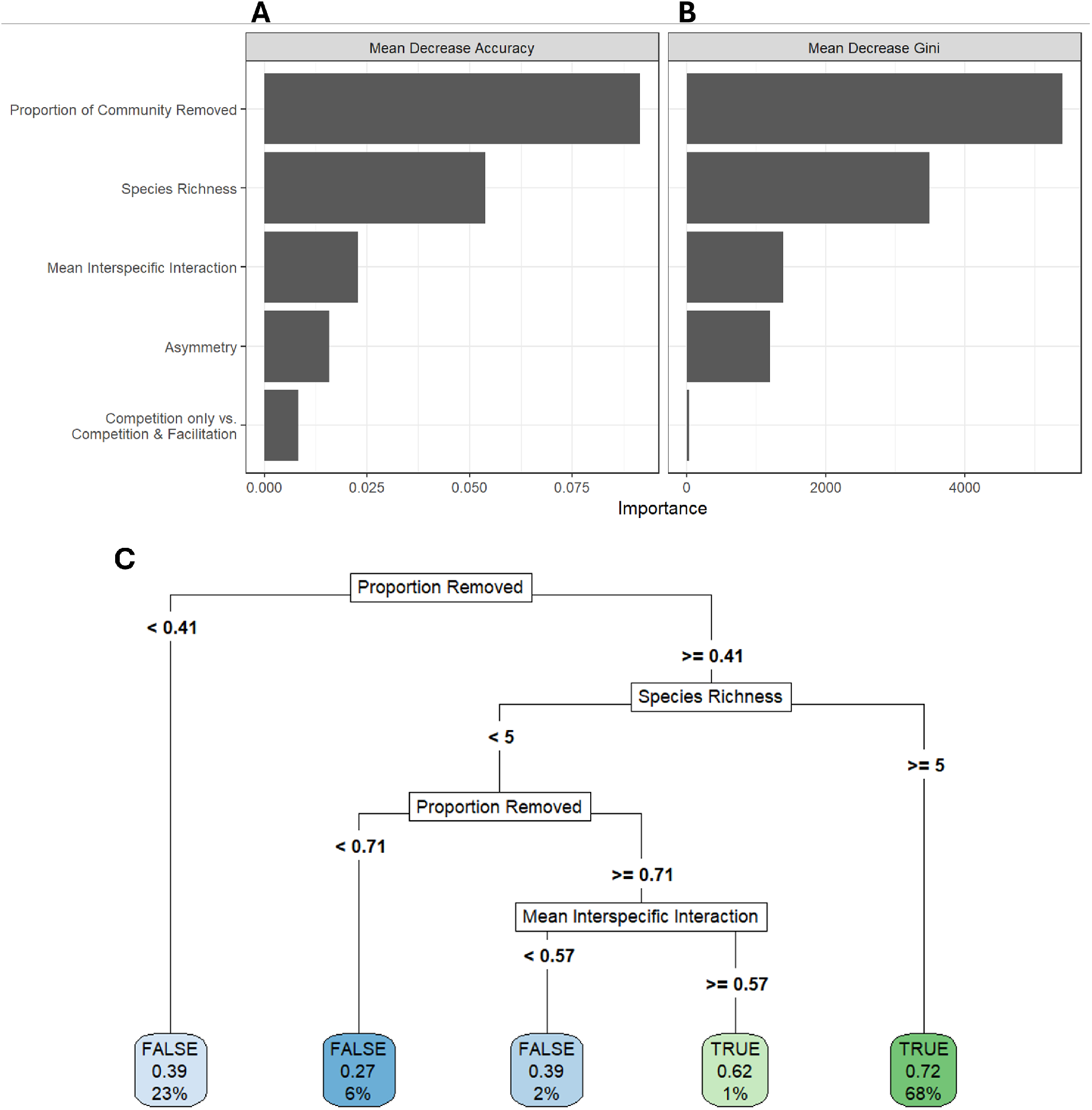
Proportion of community in extinction sequence and species richness primarily contribute to variance and likelihood of inverse priority effects. (A, B) The vertical axis depicts predictors in the model while the horizontal axis depicts the importance values of respective predictors. Panel A shows mean decrease in accuracy while B shows mean decrease in Gini. (C) Recursive partitioning tree for inverse priority effects. Internal nodes represent the predictor used for each split, and edge labels denote the threshold defining the partition. Terminal nodes indicate the predicted outcome (TRUE or FALSE), the estimated probability of an inverse priority effect for observations in that node, and the percentage of observations assigned to that node.

## Discussion

Here, we provide quantitative support that asymmetry in competitive interactions can contribute to inverse priority effects, a leading hypothesis posed in earlier studies (Torres et al., 2024a). In three-species competitive communities, those exhibiting at least one inverse priority effect consistently show higher levels of interaction asymmetry. Moreover, three-species competitive communities with multiple possible inverse priority effects tend to be more asymmetric (Appendix E). The role of asymmetry is further evident in higher-diversity communities: at low levels of asymmetry, inverse priority effects are rare, but even modest increases in asymmetry lead to a rapid increase in their probability, which eventually saturates at higher values. In general, our results demonstrate that, as hypothesized, asymmetric species interactions are a key driver of inverse priority effects and, more generally, complex community dynamics (Ranjan et al., 2024; Torres et al., 2024a; Calleja-Solanas et al., 2026).

We find that communities with stronger average competitive interactions are more prone to inverse priority effects. Previous studies have demonstrated that species removal is more likely to trigger secondary extinctions in communities where species have relatively strong interspecific competition (Fowler, 2010; Bewick et al., 2013; Sanders et al., 2015; Donohue et al., 2017). Because secondary extinctions are a key mechanism underlying inverse priority effects, we would therefore expect stronger competitive interactions to increase the likelihood of inverse priority effects, as observed in our analyses. Importantly, however, inverse priority effects can also arise in communities without strong average competition when interaction asymmetry is sufficiently high (Figure 4C, 5C), consistent with interaction asymmetry being another mechanism that can promote secondary extinctions (Fowler, 2013). Thus, competition and interaction asymmetry provide distinct yet interacting mechanisms through which species removal can generate secondary extinctions contributing to inverse priority effects.

In contrast, highly facilitative systems generate virtually no inverse priority effects. The statistical model is highly uncertain in this region of parameter space because strongly facilitative systems rarely exhibit bounded dynamics across all species combinations and therefore are uncommon in our study (Figure 5B). This pattern arises in part from our assumption that all species are capable of persisting in isolation i.e., facilitation is facultative rather than obligate. Under this assumption, strongly facilitative interactions combined with single-species persistence eliminate contingent coexistence, thereby preventing secondary extinctions and, consequently, inverse priority effects (Mierczyński and Schreiber, 2002; Barabás et al., 2014). Relaxing the assumption of guaranteed single-species persistence may increase the range of inverse priority effects in strongly facilitative systems. Additionally, incorporating saturating per-capita facilitative effects can decrease the likelihood of unbounded dynamics, potentially allowing for more coexisting, facilitative communities in which to study inverse priority effects (Buche et al., 2025).

Prior theoretical work on inverse priority effects has focused primarily on three-species competitive communities (Torres et al., 2024a). Relaxing this assumption, we find that both species richness and the fraction of species removed in an extinction sequence strongly increase the probability of observing an inverse priority effect. Our results parallel those of Case (1990, 1991), who invaded randomly constructed Lotka-Volterra competition communities. Richer and more strongly interacting communities repelled invaders more often, but successful invasions caused more resident extinctions. The mean number of residents lost per invasion (Case, 1990) and per replacement (Case, 1991) increased superlinearly with species richness and mean interaction strength. Inverse priority effects are the removal counterpart of this fragility. These patterns also echo May (1972), finding that random communities are less likely to be stable as richness and interaction strength increase, a conclusion refined by Allesina and Tang (2012).

Due to computational constraints, we restrict our analysis to communities of up to six and seven species; however, the relationships we observe are likely to change beyond this range. For example, within the richness values considered here, the probability of an inverse priority effect increases approximately linearly with species richness. We hypothesize that, as species richness continues to increase, this relationship will eventually saturate at a relatively high probability, likely towards a probability of one. In contrast, we observe a saturating relationship between inverse priority effects and the proportion of species removed in an extinction sequence. In higher-richness systems, however, we expect this relationship to become more hump-shaped, with intermediate sequence lengths maximizing the probability of inverse priority effects while very long sequences reduce it. We predict this because increasing the number of removals reduces the number of feasible coexisting communities that can serve as endpoints of disassembly, whereas intermediate extinction lengths maximize the number of possible intermediate disassembly pathways. Furthermore, including all species in an extinction sequence has no chance of an inverse priority effect, suggesting a nonmonotonic relationship between the length of the extinction sequence and the probability of an inverse priority effect. In our systems with both competition and facilitation, we partially observe this emerging pattern (Figure 5 & Appendix H).

We find that most of our qualitative results are robust to adjustments in our assumptions including the sampling distribution of interspecific interactions, definitions of coexistence for the full community and its subcommunities, and varying intraspecific competition (Appendix C, D). However, some nuances differ. For instance, when examining predictors of inverse priority effects in more speciose communities under local stability instead of permanence, we find that the probability of an inverse priority effect increases and saturates at intermediate levels asymmetry but then increases again (Appendix C). However, this pattern was not observed in systems containing both competition and facilitation (Appendix D). Several mechanisms could explain these differences. First, coexistence states that are locally but not globally attracting may have different relationships between predictors and inverse priority effects, as observed in the three-species competitive analyses with interaction asymmetry (Appendix I). Second, requiring local stability excludes coexistence states such as stable limit cycles or chaotic attractors. Excluding these dynamical behaviors may therefore alter the relationship between predictors, such as interaction asymmetry, and inverse priority effects. Finally, the observed pattern may arise under our requirement for permanence but remain undetected because relatively few communities satisfying permanence exhibit such large values of asymmetry. Regardless of the underlying mechanism, these contrasting results highlight that the stability of coexistence states can influence the drivers of inverse priority effects and the community disassembly process more generally.

## Limitations & Future Directions

We primarily use community-level metrics to quantify the prevalence of inverse priority effects. Thus, a key future direction is to better resolve how species-level attributes shape these outcomes. Torres et al. (2024a) addressed this issue using niche and fitness differences between species pairs within the system. However, in more species-rich communities and in the presence of non-competitive interactions, defining niche and fitness differences is challenging (Godwin et al., 2020), motivating the use of alternative metrics (though, see Spaak et al. (2021)). Promising candidates include measures based on the relative strength of species interactions or the asymmetry of interactions involving the species removed relative to the rest of the community. Moreover, rather than computing these quantities solely at the level of the full community, they could be evaluated along disassembly pathways. This would allow for a more dynamic characterization of how interaction structure changes through extinction sequences and may help determine whether metrics such as asymmetry or mean interaction strength along disassembly trajectories can predict the emergence of inverse priority effects.

Here, we assume that following the removal of a species, the system settles to the corresponding disassembled community before the next extinction occurs. While this assumption simplifies the analysis, it excludes cases in which subsequent removals occur during transient dynamics (Hastings et al., 2018; Grainger et al., 2025). For example, in Figure 2, an inverse priority effect arises because species 2 excludes species 1 following the removal of species 3. However, if species 2 is removed sufficiently quickly after species 3, species 1 may persist by avoiding competitive exclusion, thereby eliminating the inverse priority effect. Although the community disassembly graph partially accounts for timing effects by allowing multiple outgoing edges from a given coexistence state when extinction timing influences the disassembled community (Brennan and Schreiber, 2026), it still assumes that the system is initially at some community composition that can successfully coexist before each removal. Future work that explicitly examines how inverse priority effects can be both eliminated and generated when species removals occur during transient dynamics would provide a more nuanced understanding of these dynamics. Another interesting extension would be to develop community disassembly graphs that explicitly incorporate simultaneous extinction events, enabling direct comparison between sequential and concurrent removals. This would be a relatively simple extension of the conditions provided in Brennan and Schreiber (2026). Such an approach could clarify how the timing of extinction shapes inverse priority effects and would closely reflect the experimental tests and observations of these phenomena (Collins et al., 2009; Torres et al., 2024b).

As in previous studies, we assume that all species within a community have identical, positive intrinsic growth rates (Jiang and Liang, 2020; Torres et al., 2024a). Although this allows us to precisely disentangle the impacts of species interactions in driving inverse priority effects, it also restricts the possible dynamics in the Lotka-Volterra model. Future studies that loosen this assumption to allow variable positive intrinsic growth rates can help elucidate how variance in both species interactions and intrinsic growth rates can impact the prevalence of inverse priority effects. Furthermore, a natural next step would be allowing for both positive and negative intrinsic growth rates, as in multi-trophic level systems such as in predator-prey food webs (Borrvall and Ebenman, 2006; Emary and Evans, 2021) or plant-pollinator networks (Rohr et al., 2014; Grilli et al., 2017). We hypothesize that multi-trophic systems that have coexistence mediated by indirect effects such as keystone predation will be more likely to have inverse priority effects than those that do not. While direct trophic dependencies that lead to secondary extinction (e.g., loss of prey resulting in predator extinction) may contribute to inverse priority effects, we expect these effects to be weaker than those driven by indirect interactions spanning trophic levels.

Here, we model community dynamics using the Lotka-Volterra model not only because of its keystone role in community ecology but also for analytical and numerical tractability. Despite its utility, Lotka-Volterra models can neglect ecological processes that are possibly important for inverse priority effects. For example, ecosystem engineers, which often play key roles in communities, can modify abiotic conditions with indirect consequences for other species (Padilla and Pugnaire, 2006; Romero et al., 2015). In a Lotka-Volterra framework, such effects are typically represented as direct interactions. As a result, extinction of an ecosystem engineer may lead to predicted secondary extinctions of species that depend on its modification of abiotic conditions. However, in some systems, persistence may depend only on the presence of a suitably conditioned environment, rather than continued interaction with the engineer itself. In such cases, even if species cannot establish in the absence of the ecosystem engineer, they may persist following its extinction provided that the environmental state remains altered (Reynolds et al., 2017; Albertson et al., 2024). Such dynamics can therefore alter patterns of secondary extinction and, consequently, inverse priority effects. Although the community disassembly graph framework can account for these scenarios (Brennan and Schreiber, 2026), doing so would require a different model of community dynamics and be more mathematically and computationally challenging. Nevertheless, such extensions represent an important direction for future work toward establishing general principles of inverse priority effects, particularly for inference in empirical systems.

## Conclusion

We show that both interaction asymmetry and interaction strength contribute to inverse priority effects, quantitatively supporting previous hypotheses. We further demonstrate that inverse priority effects become more prevalent with increasing species richness and likely vary nonmonotonically with the fraction of species removed. Our methodology provides a framework for identifying when inverse priority effects arise, and our results reveal the ecological properties that promote them. The high prevalence of inverse priority effects in communities of up to six or seven species suggests that they might be common in empirical communities, particularly those characterized by high species richness, strong competition, and asymmetric interactions, thus warranting further investigation of inverse priority effects and complex dynamics of community disassembly more generally.

## Acknowledgments

We thank Marissa Baskett and John Stachowicz for feedback on an early draft. We thank Raissa D’Souza, Alan Hastings, Marcel Holyoak, and participants of the Hastings-Schreiber joint lab meeting for constructive feedback throughout the development of this project. JB acknowledges support from the National Science Foundation Graduate Research Fellowship program under Grant No. DGE-2439024.

~~~
File: main.tex
Encoding: ascii
Sum count: 8394
Words in text: 6700
Words in headers: 187
Words outside text (captions, etc.): 1358
Number of headers: 35
Number of floats/tables/figures: 22
Number of math inlines: 140
Number of math displayed: 9
Subcounts:
    text+headers+captions (#headers/#floats/#inlines/#displayed)
    28+12+0 (1/0/1/0) _top_
    54+1+0 (1/0/0/0) Section: Acknowledgments
    39+2+0 (1/0/0/0) Section: Author Contributions
    182+1+0 (1/0/0/0) Section: Abstract
    686+1+0 (1/0/0/0) Section: Introduction
    0+1+0 (1/0/0/0) Section: Methods
    762+4+141 (1/1/36/3) Subsection: Three species competitive communities
    784+4+110 (1/1/47/1) Subsection: More speciose competitive communities
    525+6+0 (1/0/4/0) Subsection: Speciose communities with competition and
    0+1+0 (1/0/0/0) Section: Results
    96+4+89 (1/1/1/0) Subsection: Three species competitive communities
    175+4+110 (1/1/0/0) Subsection: More speciose, competitive communities
    323+6+235 (1/2/1/0) Subsection: Speciose communities with competition a
    947+1+0 (1/0/0/0) Section: Discussion
    773+3+0 (1/0/6/0) Section: Limitations \& Future Directions
    124+1+0 (1/0/0/0) Section: Conclusion
    52+7+0 (1/0/8/5) Section: Lotka Volterra change of coordinates and
    28+6+0 (1/0/3/0) Section: Parameters for dynamical classes of interest
    86+5+0 (1/0/2/0) Section: Competitive analyses robustness to assumption
    71+6+38 (1/1/3/0) Subsection: Different sampling distributions of inter
    135+8+47 (1/1/2/0) Subsection: Local instead of global stability of coe
    80+5+38 (1/1/0/0) Subsection: Requiring permanence of all subcommunitie
    50+3+40 (1/1/4/0) Subsection: Varying intraspecific competition
    129+4+40 (1/1/4/0) Subsection: Allowing cyclic invasion graphs
    42+6+43 (1/1/1/0) Subsection: Different sampling distributions of inter
    53+8+45 (1/1/0/0) Subsection: Local instead of global stability of coex
    31+5+41 (1/1/0/0) Subsection: Requiring permanence of all subcommunitie
    49+3+40 (1/1/3/0) Subsection: Varying intraspecific competition
    89+7+0 (1/0/2/0) Section: Competitive and facilitative analyses robustn
    111+4+45 (1/1/4/0) Subsection: Allowing cyclic invasion graphs
    0+10+34 (1/1/3/0) Section: All possible inverse priority effects in thr
    109+7+19 (1/1/1/0) Section: Asymmetry of each possible three species co
    0+15+28 (1/1/0/0) Section: Empirical values for combinations of species
    0+18+34 (1/1/0/0) Section: Empirical values for combinations of species
    87+8+141 (1/2/4/0) Section: Locally stable rather than permanent three
File: output.bbl
Encoding: utf8
Sum count: 0
Words in text: 0
Words in headers: 0
Words outside text (captions, etc.): 0
Number of headers: 0
Number of floats/tables/figures: 0
Number of math inlines: 0
Number of math displayed: 0
Total
Sum count: 8394
Words in text: 6700
Words in headers: 187
Words outside text (captions, etc.): 1358
Number of headers: 35
Number of floats/tables/figures: 22
Number of math inlines: 140
Number of math displayed: 9
Files: 2
Subcounts:
     text+headers+captions (#headers/#floats/#inlines/#displayed)
     6700+187+1358 (35/22/140/9) File(s) total: main.tex
     0+0+0 (0/0/0/0) File(s) total: output.bbl
(errors:5)
~~~

## A Lotka Volterra change of coordinates and nondimensionalization

Let

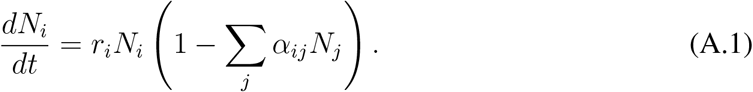

Let *x*_*i*_ = *α*_*ii*_*N*_*i*_. Then, we have 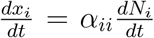. Performing this change of coordinates, we have

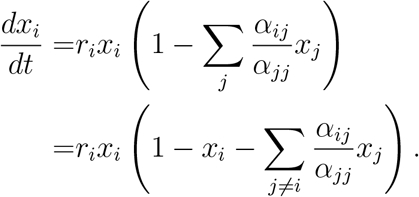

Let 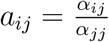 . Then, we have

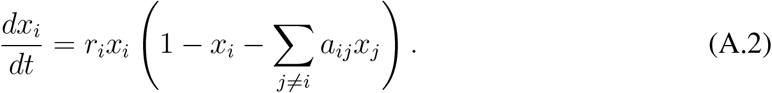

Let all species *i ∈ {*1, …, *n}* have an equal intrinsic growth rate of *r*. We then have

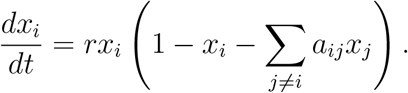

Let *τ* = *rt*. By the chain rule of differentation, we have that 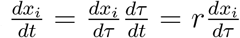 . Dividing both sides of the equation by *r*, we arrive at our final model:

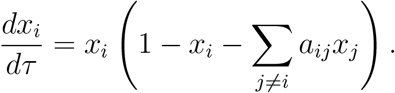

## B Parameters for dynamical classes of interest

**Table B.1:**
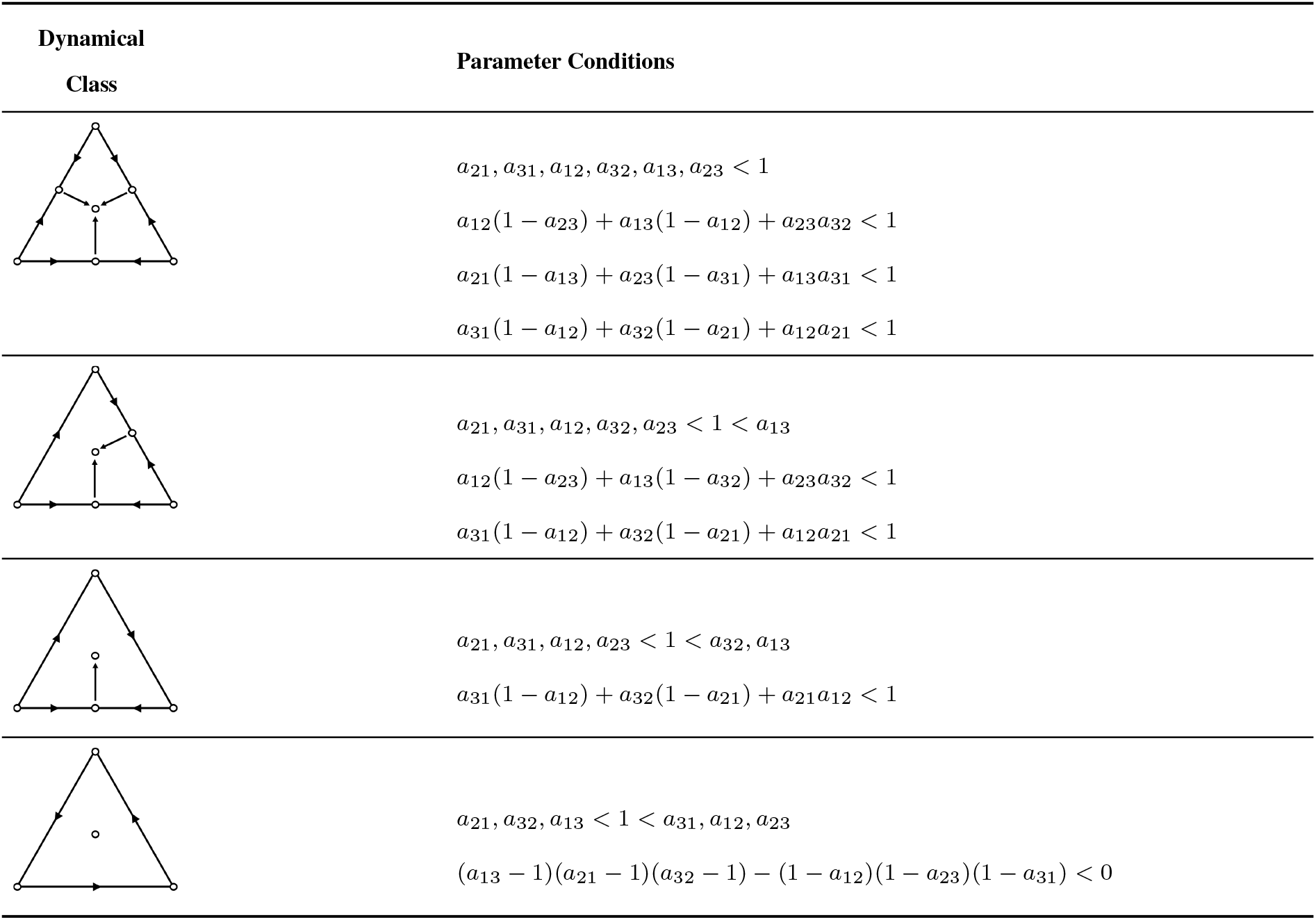
Parameter conditions for each dynamical class under Equation 3. Parameter conditions derived in Jiang and Liang (2020). Bottom left node denotes species 1, bottom right node denotes species 2, and top node denotes species 3.

| Dynamical Class | Parameter Conditions |
| --- | --- |
| | $a_{21}, a_{31}, a_{12}, a_{32}, a_{13}, a_{23} < 1$<br>$a_{12}(1 - a_{23}) + a_{13}(1 - a_{12}) + a_{23}a_{32} < 1$<br>$a_{21}(1 - a_{13}) + a_{23}(1 - a_{31}) + a_{13}a_{31} < 1$<br>$a_{31}(1 - a_{12}) + a_{32}(1 - a_{21}) + a_{12}a_{21} < 1$ |
| | $a_{21}, a_{31}, a_{12}, a_{32}, a_{23} < 1 < a_{13}$<br>$a_{12}(1 - a_{23}) + a_{13}(1 - a_{32}) + a_{23}a_{32} < 1$<br>$a_{31}(1 - a_{12}) + a_{32}(1 - a_{21}) + a_{12}a_{21} < 1$ |
| | $a_{21}, a_{31}, a_{12}, a_{23} < 1 < a_{32}, a_{13}$<br>$a_{31}(1 - a_{12}) + a_{32}(1 - a_{21}) + a_{21}a_{12} < 1$ |
| | $a_{21}, a_{32}, a_{13} < 1 < a_{31}, a_{12}, a_{23}$<br>$(a_{13} - 1)(a_{21} - 1)(a_{32} - 1) - (1 - a_{12})(1 - a_{23})(1 - a_{31}) < 0$ |

## C Competitive analyses robustness to assumptions

Here, we present our results when we change various assumptions in our analyses of speciose, competitive communities. The order of assumptions tested are as follows: (1) distributional assumptions of interspecific competition coefficients, (2) local instead of global stability of the full coexistence state, (3) permanence rather than feasibility of subcommunities, (4) variation in intraspecific competition, and (5) positive invasion growth rates of species *i* at all *−i* communities of the full coexistence state rather than permanence. All statistical models share the same form as in the main text.

### C.1 Different sampling distributions of interspecific competition

First, we test the distributional assumptions for the parameter draws by considering three alternatives: lognormal, uniform and exponential distributions. For the lognormal case, we sample 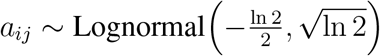, which yields, after exponentiation, a distribution with mean and variance equal to one. For the uniform case, we follow the parameter range used in Case (1990), drawing *a*_*ij*_ *∼ U* (0, 1.6). For the exponential case, *a*_*ij*_ *∼* Exp(1). Across all alternatives, we find that the qualitative dynamics remain unchanged relative to those shown in Figure 4.

**Figure C.1.**
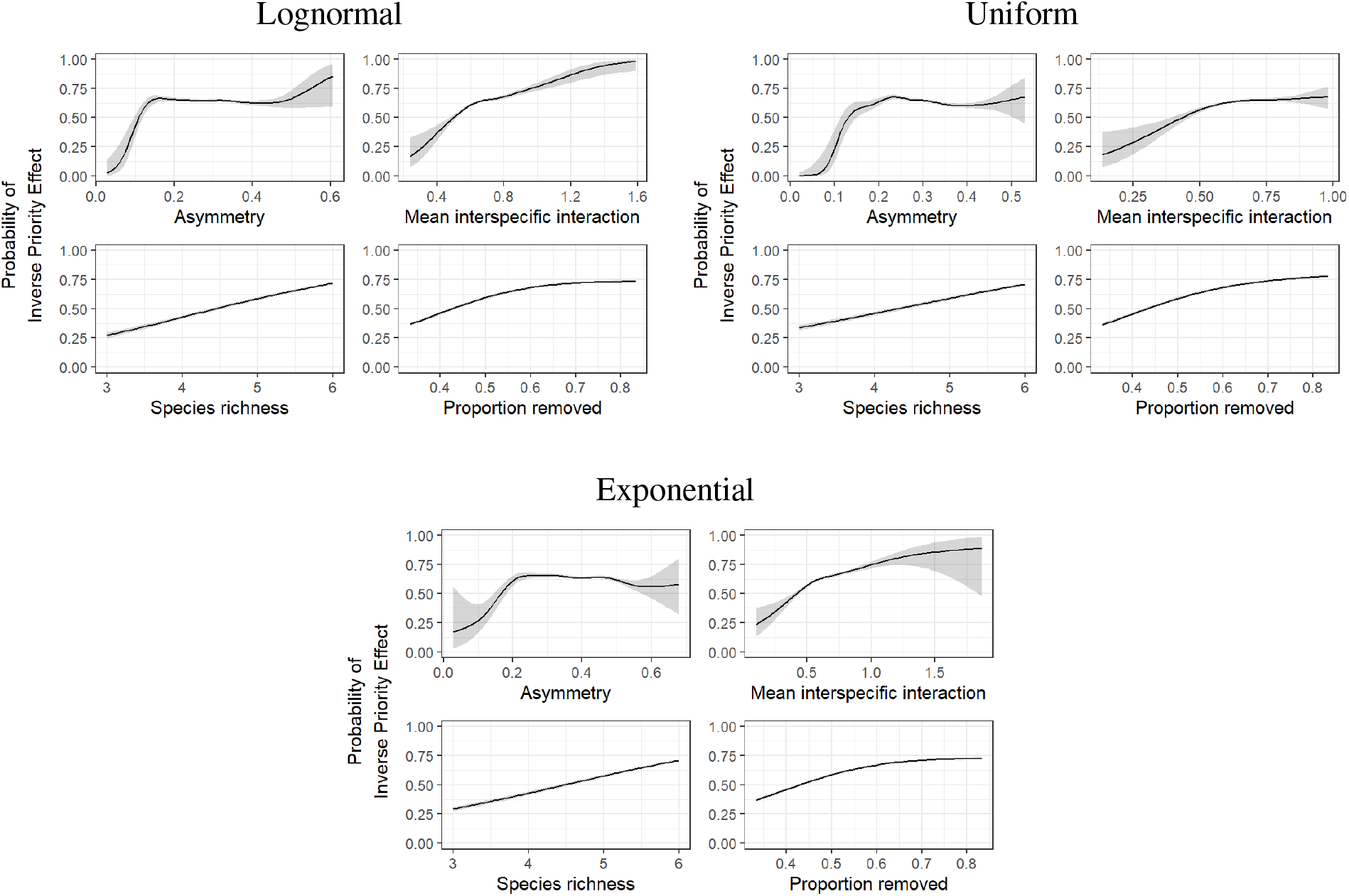
Different distributions give same qualitative results for competitive communities. Plotted are the conditional predicted probabilities of an inverse priority effect across the range of each predictor, with all other predictors held at their observed means.

### C.2 Local instead of global stability of coexistence state

Next, we assess how our results may differ if we allow the full coexistence state to coexist under a locally stable equilibrium rather than a global attractor. Given the arbitrarily complex dynamics Lotka-Volterra can provide when there are more than two species, a set of species may coexist under a locally stable equilibrium yet not be a global attractor for the full system. We find that when we only check for local stability of the full coexistence state, we generally see the same qualitative trends for each predictor of interest. The one exception is that, for asymmetry, we see an initial saturation past *∼* 0.2 until *∼* 0.5, when the probability of inverse priority effect begins increasing again with large asymmetry values. Note that our analyses requiring permanence of the fully community do not reach asymmetry values that large.

**Figure C.2:**
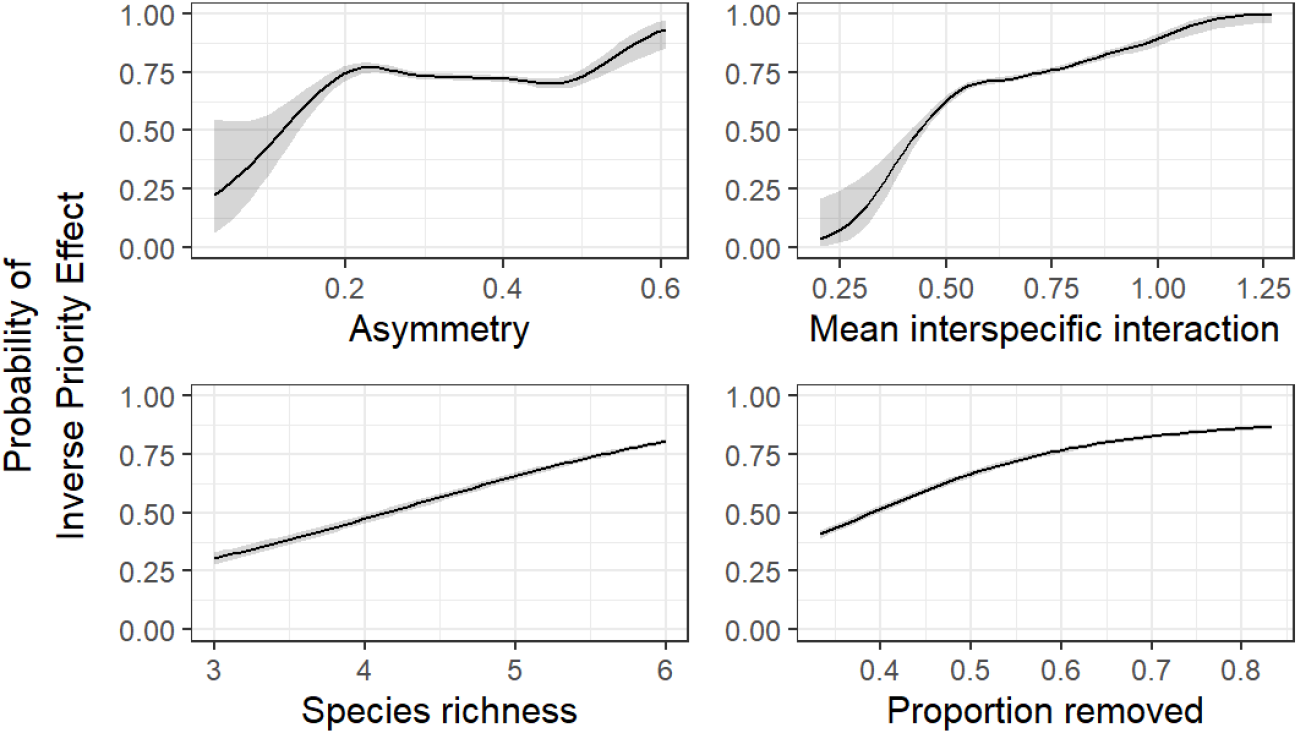
Local rather than global stability of the full coexistence state does not change qualitative dynamics for most predictors in competitive communities. Plotted are the conditional predicted probabilities of an inverse priority effect across the range of each predictor, with all other predictors held at their observed means.

### C.3 Requiring permanence of all subcommunities

Next, we test whether requiring permanence for all subcommunities alters the qualitative results. In some cases, a subcommunity may have a positive equilibrium without being feasible (e.g. in the presence of bistability). In this case, the community disassembly graph would suggest a possible disassembly trajectory toward that community, even though it is effectively unreachable in simulations. Imposing permanence as a requirement for all subcommunities eliminates such cases. Despite this more restrictive assumption, we find that the qualitative dynamics remain unchanged.

**Figure C.3:**
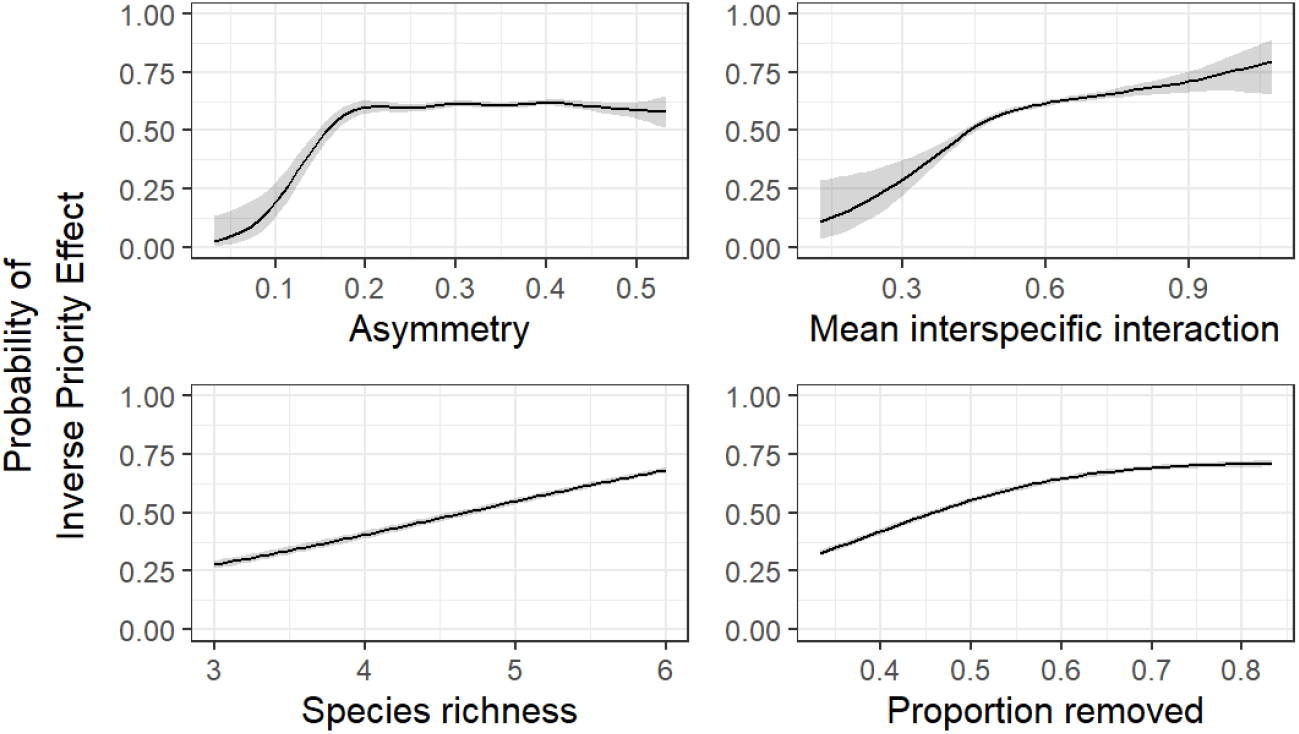
Requiring permanence of subcommunities does not change qualitative dynamics for competitive communities. Plotted are the conditional predicted probabilities of an inverse priority effect across the range of each predictor, with all other predictors held at their observed means.

### C.4 Varying intraspecific competition

Next, we test if qualitative findings are consistent even with variance in intraspecific competition. To this end, we sample parameters *a*_*ij*_ from Equation 1 such that *a*_*ij*_ = |*Z*_*ij*_|, *Z*_*ij*_ *∼ N* (0, 1) for all *i, j* including *i* = *j*. We find that qualitative relationships between predictors and inverse priority effects mostly remain similar even when allowing for variation in intraspecific competition (Figure C.4).

**Figure C.4:**
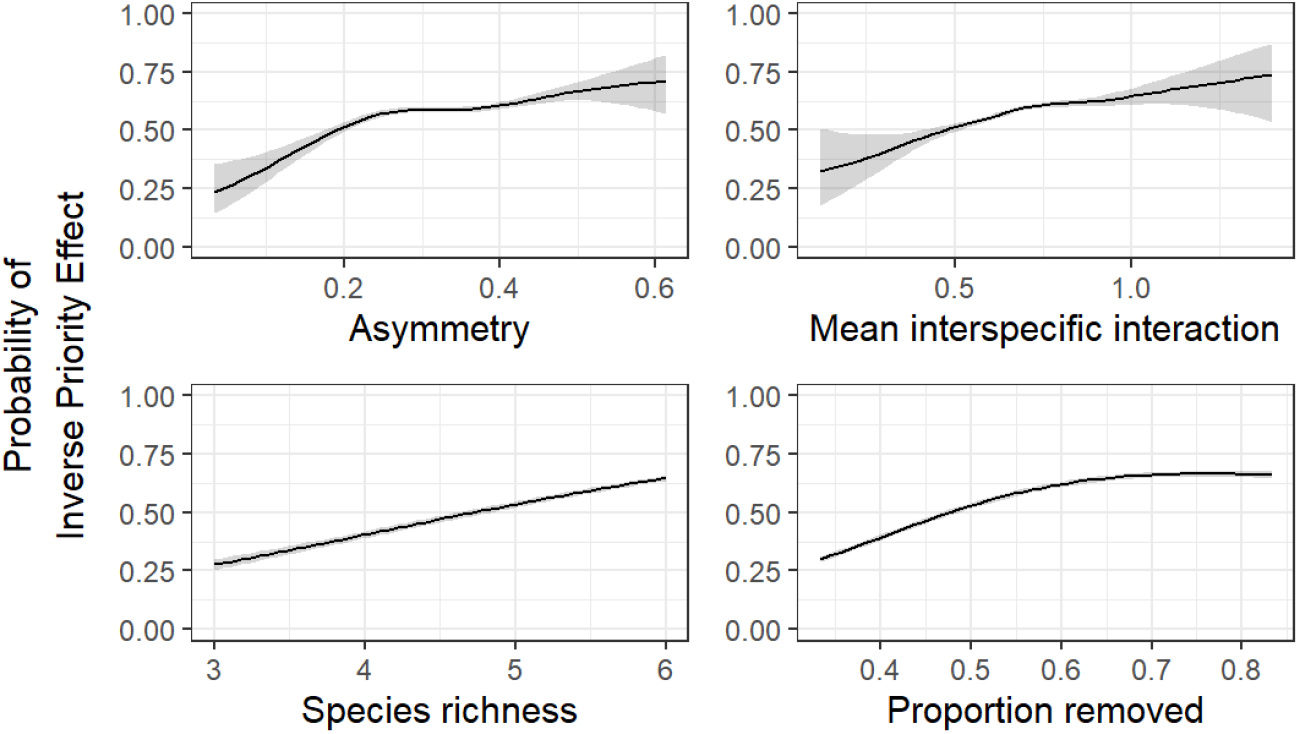
Varying intraspecific competition does not change qualitative dynamics for communities with competition and facilitation. Plotted are the conditional predicted probabilities of an inverse priority effect across the range of each predictor, with all other predictors held at their observed means.

### C.5 Allowing cyclic invasion graphs

Lastly, we test if qualitative findings are consistent when we allow the full community to persist under a weaker form of persistence. In the main text, we classify coexistence of a full community when all species *i* can invade their respective *−i* communities, and there are no cycles in the invasion graph. The latter condition is generally required for this condition to be sufficient e.g., rock-paper-scissors dynamics meet the mutual invasibility condition but do not always coexist in the sense of permanence (Chesson, 2018). However, we can instead study a weaker form of persistence, requiring only the necessary condition stating that all species *i* must have a positive invasion growth rate at their *−i* community. We find that qualitative relationships between predictors and inverse priority effects are consistent when allowing cycles in the invasion graph.

**Figure C.5:**
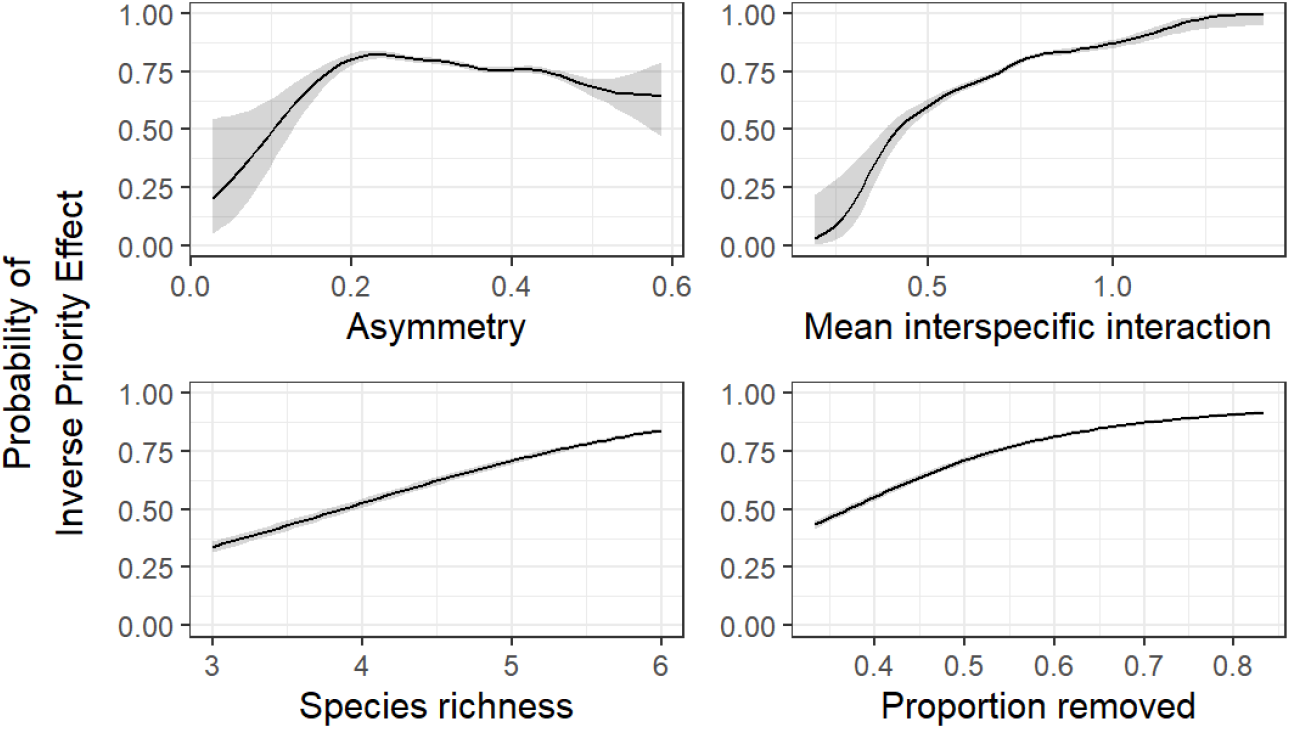
Allowing cycles in the invasion graph does not change qualitative dynamics for competitive communities. Plotted are the conditional predicted probabilities of an inverse priority effect across the range of each predictor, with all other predictors held at their observed means.

## D Competitive and facilitative analyses robustnes to assumptions

Here, we present our results when we change various assumptions in our analyses of speciose communities with competition and facilitation. The order of assumptions tested are as follows: (1) distributional assumptions of interspecific interaction coefficients, (2) local instead of global stability of the full coexistence state, (3) permanence rather than feasibility of subcommunities, (4) variation in intraspecific competition, and (5) positive invasion growth rates of species *i* at all *−i* communities of the full coexistence state rather than permanence. All statistical models share the same form as in the main text.

### D.1 Different sampling distributions of interspecific interactions

First, we test the distributional assumptions for the parameter draws by considering also a uniform distribution. We sample *a*_*ij*_ *∼ U* (*−*1.6, 1.6), similar to the parameter ranges in Case (1990). For the uniform distribution, we find that the qualitative dynamics remain unchanged relative to those shown in Figure 5.

**Figure D.1:**
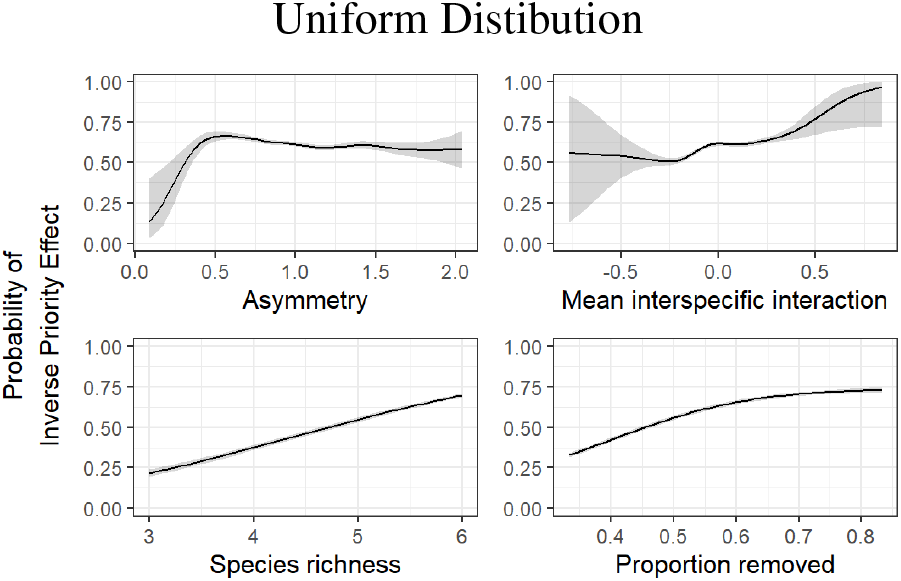
Different distributional assumption gives the same qualitative results for communities with both facilitation and competition. Plotted are the conditional predicted probabilities of an inverse priority effect across the range of each predictor, with all other predictors held at their observed means.

### D.2 Local instead of global stability of coexistence state

Next, we assess how our results may differ if we allow the full coexistence state to coexist under a locally stable equilibrium rather than a global attractor. We find that when we only check for local stability of the full coexistence state, we see the same qualitative trends for each predictor of interest.

**Figure D.2:**
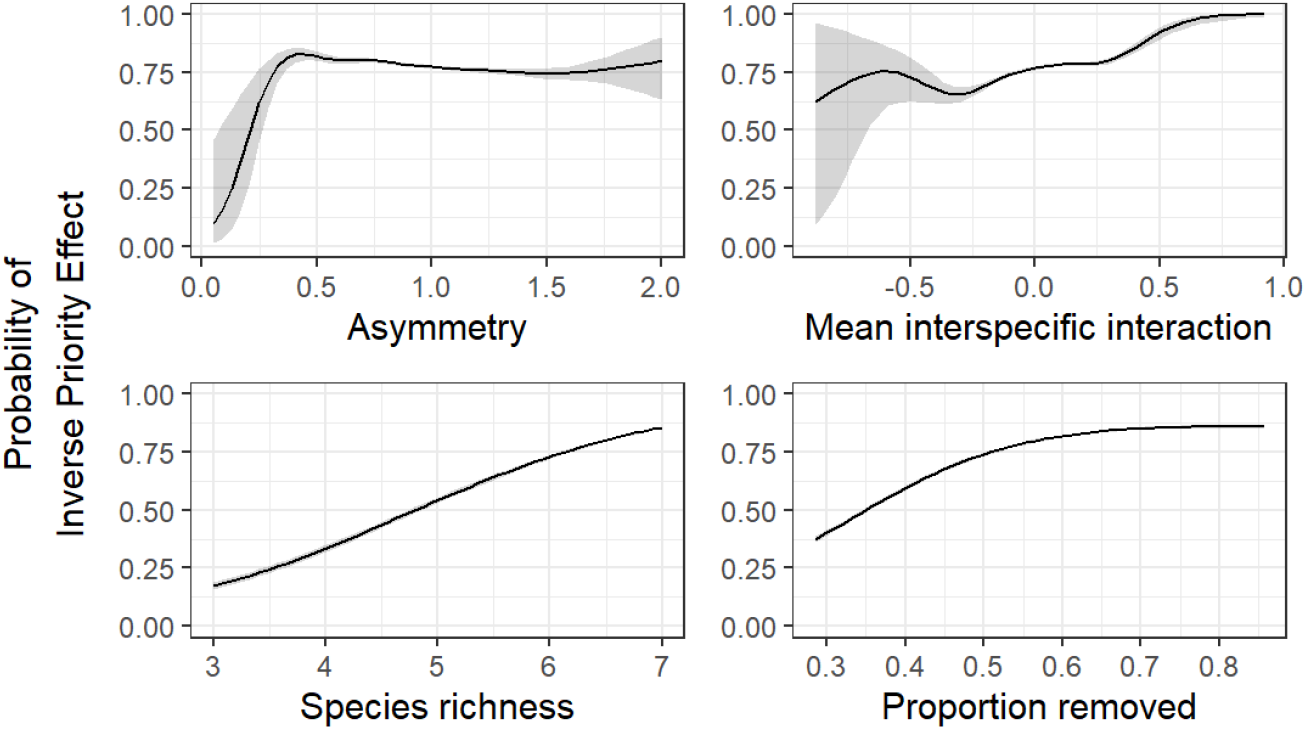
Allowing local rather than global stability of full community recovers same qualitative patterns in communities with facilitation and competition. Plotted are the conditional predicted probabilities of an inverse priority effect across the range of each predictor, with all other predictors held at their observed means.

### D.3 Requiring permanence of all subcommunities

Next, we again test whether requiring permanence for all subcommunities alters the qualitative results. Similar to the fully competitive case, requiring all subcommunities to be permanent does not change qualitative insights.

**Figure D.3:**
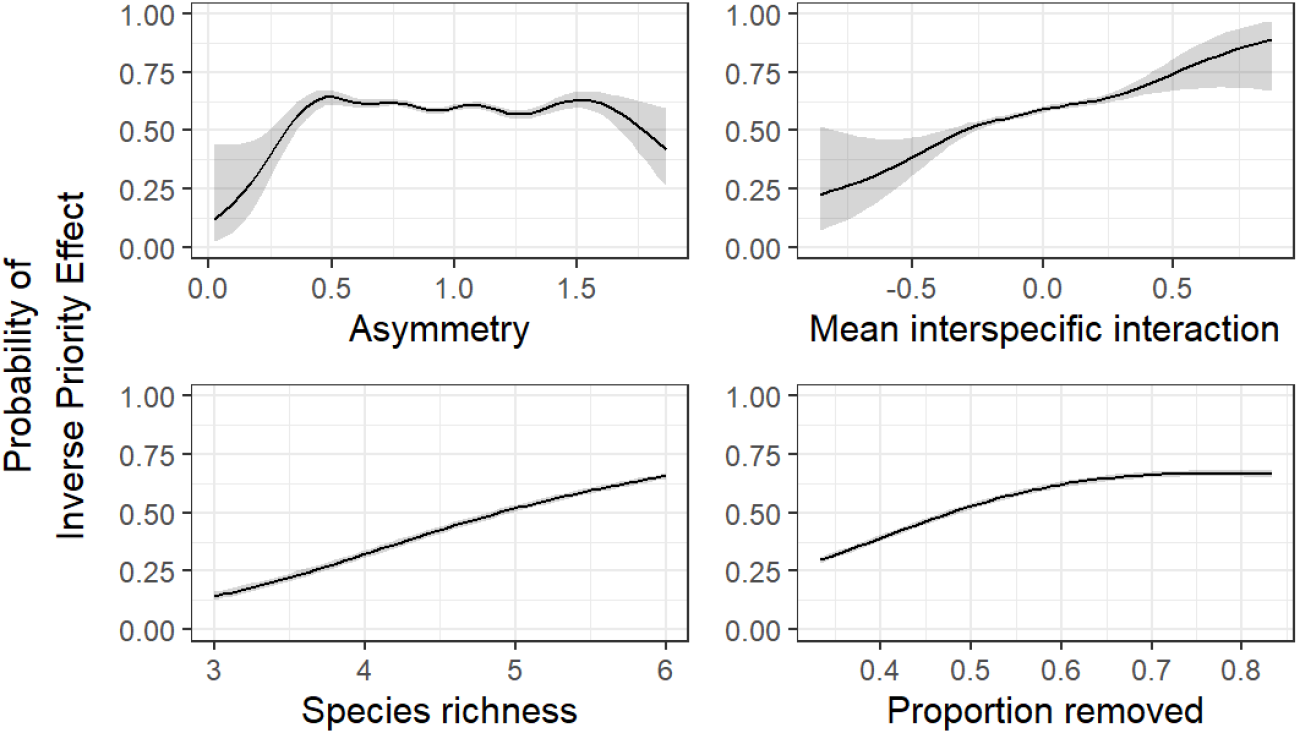
Requiring permanence of subcommunities does not change qualitative dynamics for communities with competition and facilitation. Plotted are the conditional predicted probabilities of an inverse priority effect across the range of each predictor, with all other predictors held at their observed means.

### D.4 Varying intraspecific competition

Next, we test whether qualitative findings are consistent even with variance in intraspecific competition. To this end, we sample the parameters *α*_*ij*_ from Equation 1 such that when *i≠ j, α*_*ij*_ *∼ N* (0, 1) while *a*_*ii*_ = |*Z*_*ii*_|, *Z*_*ii*_ *∼ N* (0, 1). We find that qualitative relationships between predictors and inverse priority effects remain similar even when allowing for variation in intraspecific competition (Figure D.4).

**Figure D.4:**
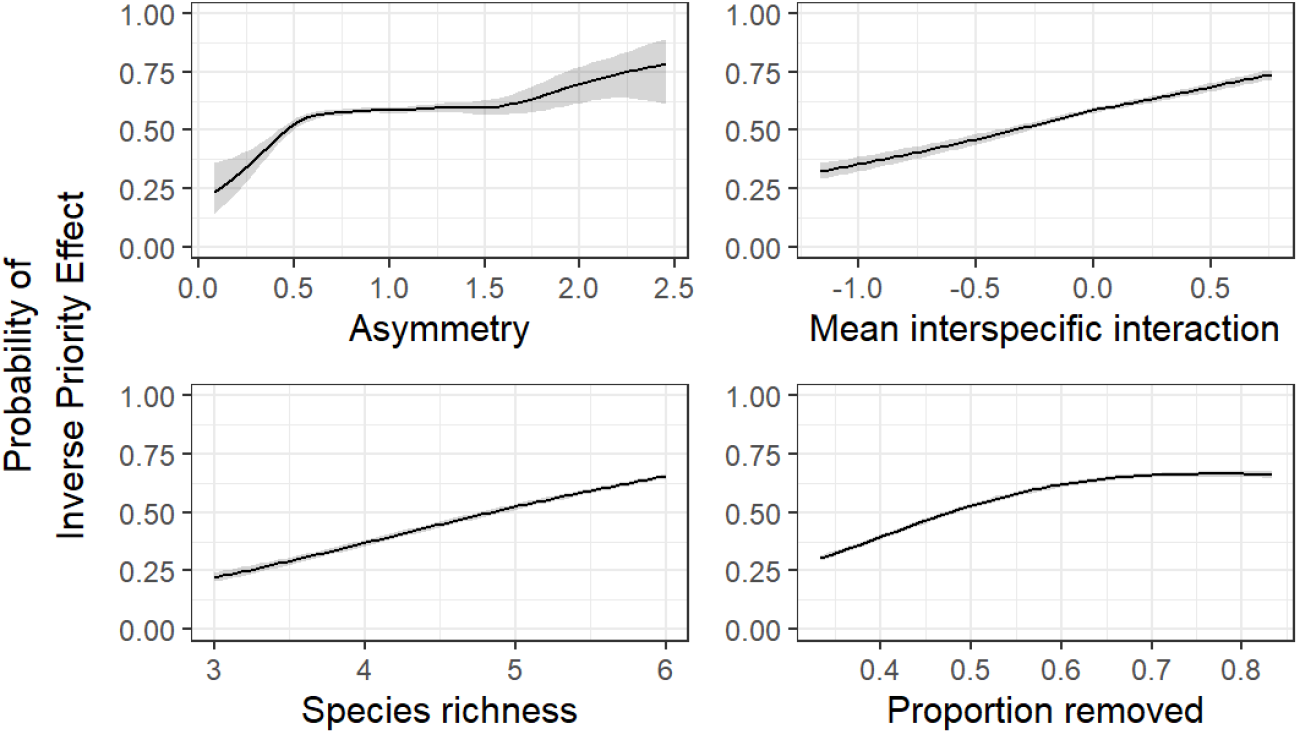
Varying intraspecific competition does not change qualitative dynamics for communities with competition and facilitation. Plotted are the conditional predicted probabilities of an inverse priority effect across the range of each predictor, with all other predictors held at their observed means.

### D.5 Allowing cyclic invasion graphs

Lastly, we test if qualitative findings are consistent when we allow the full community to persist under a weaker form of persistence. In the main text, we classify coexistence of a full community when all species *i* can invade their respective *−i* communities, and there are no cycles in the invasion graph. The latter condition is generally required for this condition to be sufficient. However, we can instead study a weaker form of persistence, requiring only the necessary condition stating that all species *i* must have a positive invasion growth rate at their *−i* community. We find that qualitative relationships between predictors and inverse priority effects are consistent when allowing cycles in the invasion graph.

**Figure D.5:**
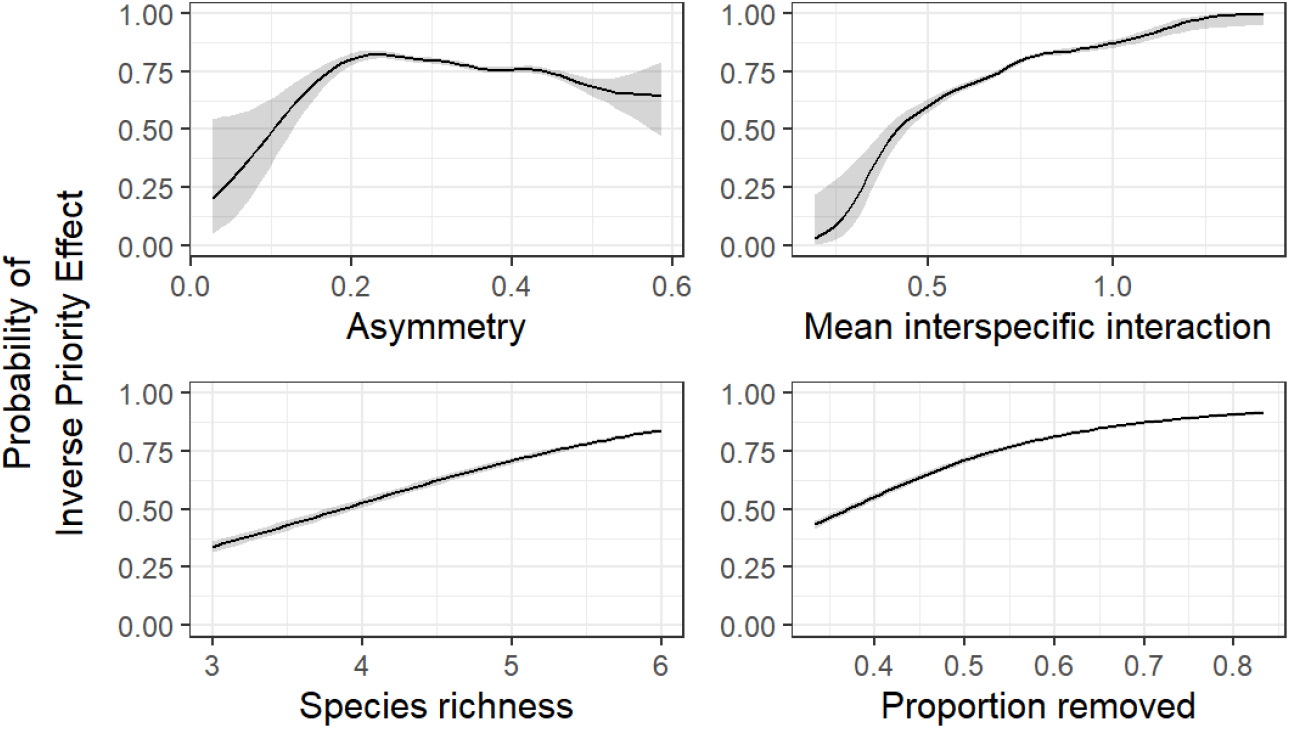
Allowing cycles in the invasion graph does not change qualitative dynamics for communities with both competitive and facilitative interactions. Plotted are the conditional predicted probabilities of an inverse priority effect across the range of each predictor, with all other predictors held at their observed means.

## E All possible inverse priority effects in three-species, competitive Lotka-Volterra models

**Figure E.1:**
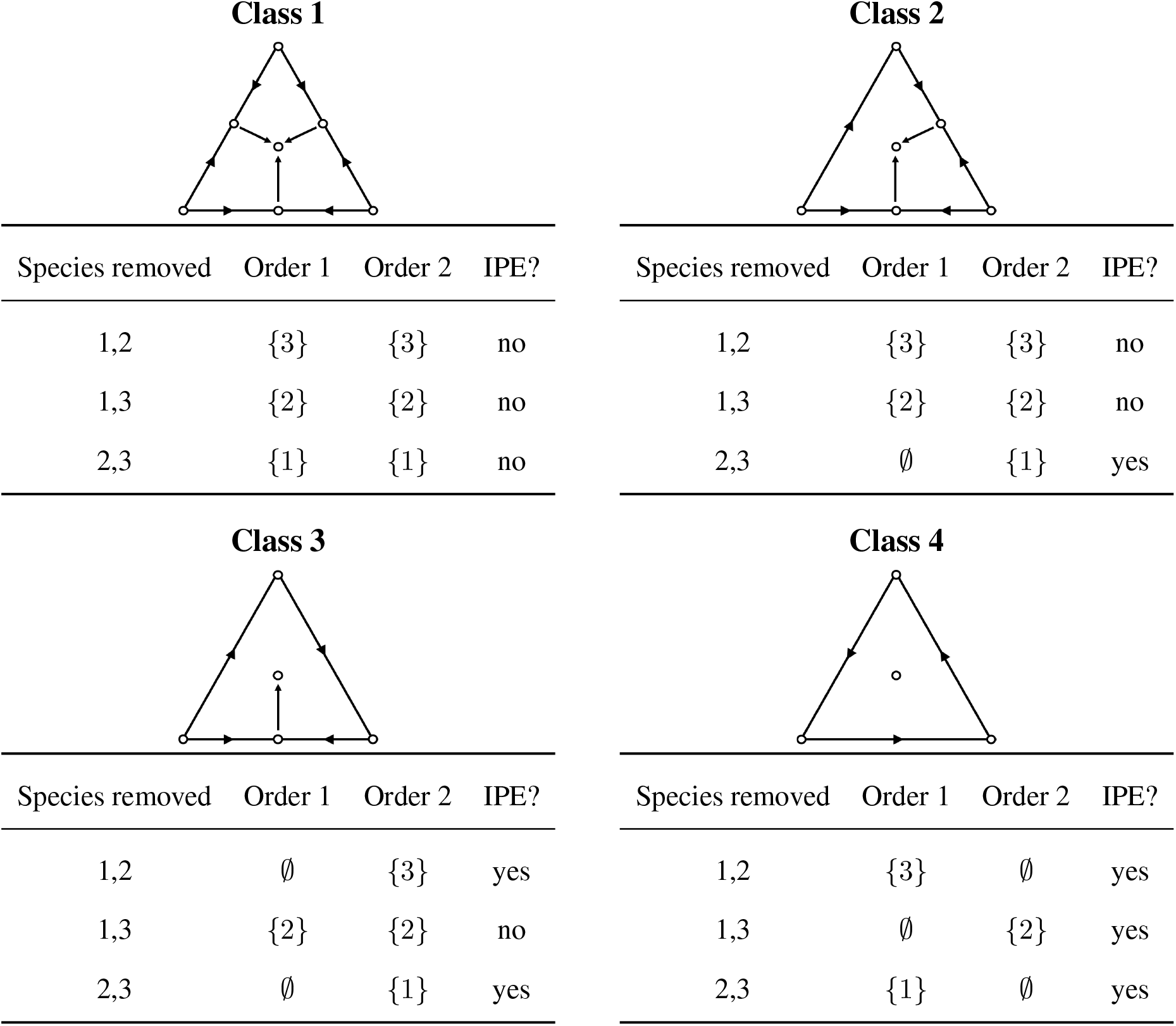
Identifying all possible inverse priority effects (IPE) across the four classes of dynamic behaviors for competitive three-species Lotka-Volterra models. Bottom left node denotes species 1, bottom right node denotes species 2, and top node denotes species 3.

## F Asymmetry of each possible three species community

Here, we demonstrate that another measure of asymmetry provides the same qualitative results in the three-species, competitive case. The alternate asymmetry metric is cyclic asymmetry, originally defined in Ranjan et al. (2024) quantifying the nonadditivity of competition between species. Here, we define a modified metric by applying the absolute value to the logarithm of the original quantity proposed in Ranjan et al. (2024), resulting in 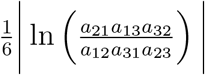. If equal to zero, there is no cyclic asymmetry in the system. As values become larger, cyclic asymmetry increases. Note that, in Figure F.1, both metrics of asymmetry provide the same qualitative results, demonstrating that the conclusion that more asymmetric interactions tend to be associated with inverse priority effects is consistent across metrics.

**Figure F.1:**
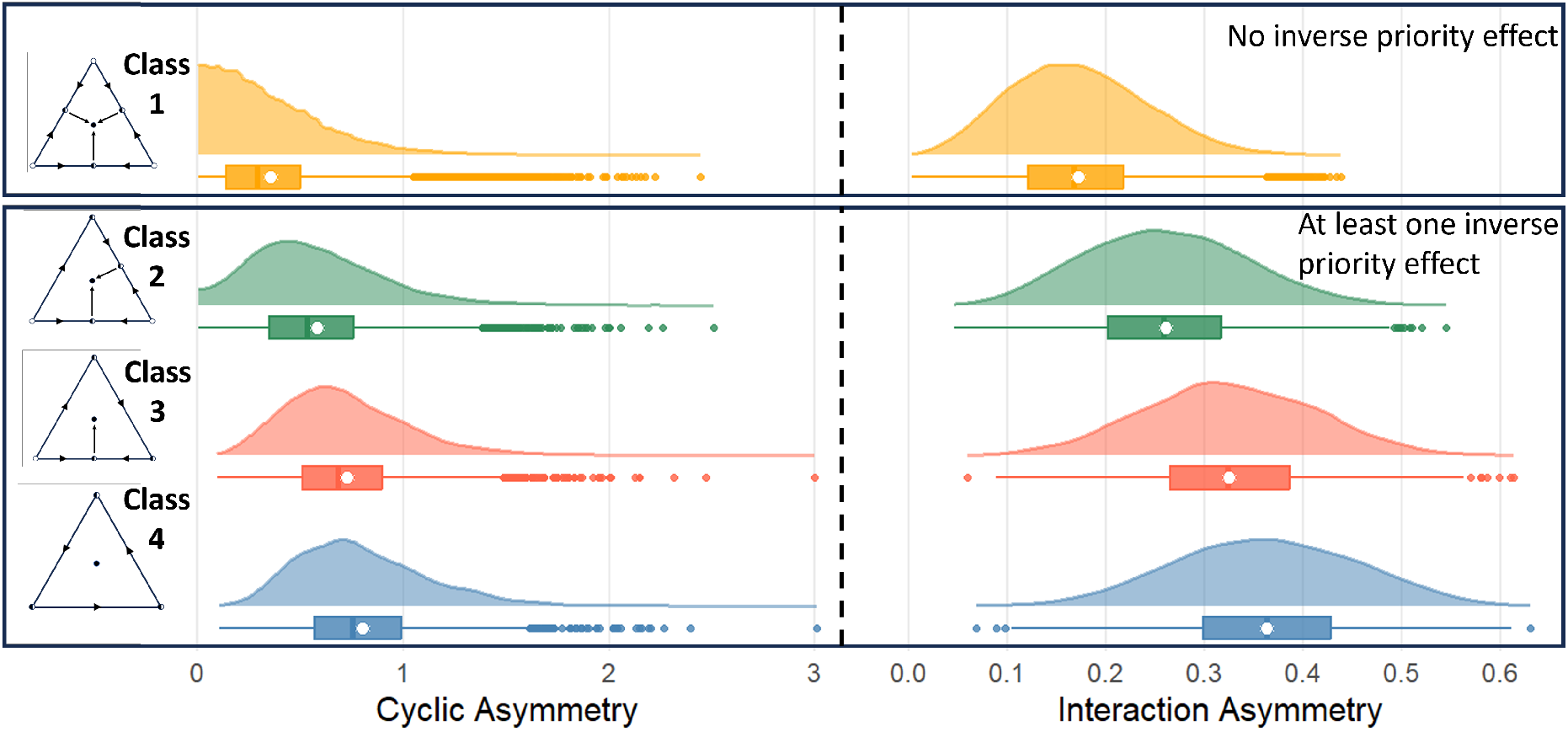
Density plots of asymmetry of simulated parameterizations across each possible class of dynamics. White circle denote the mean value.

## G Empirical values for combinations of species richness and length of extinction sequence in competitive communities

**Figure G.1:**
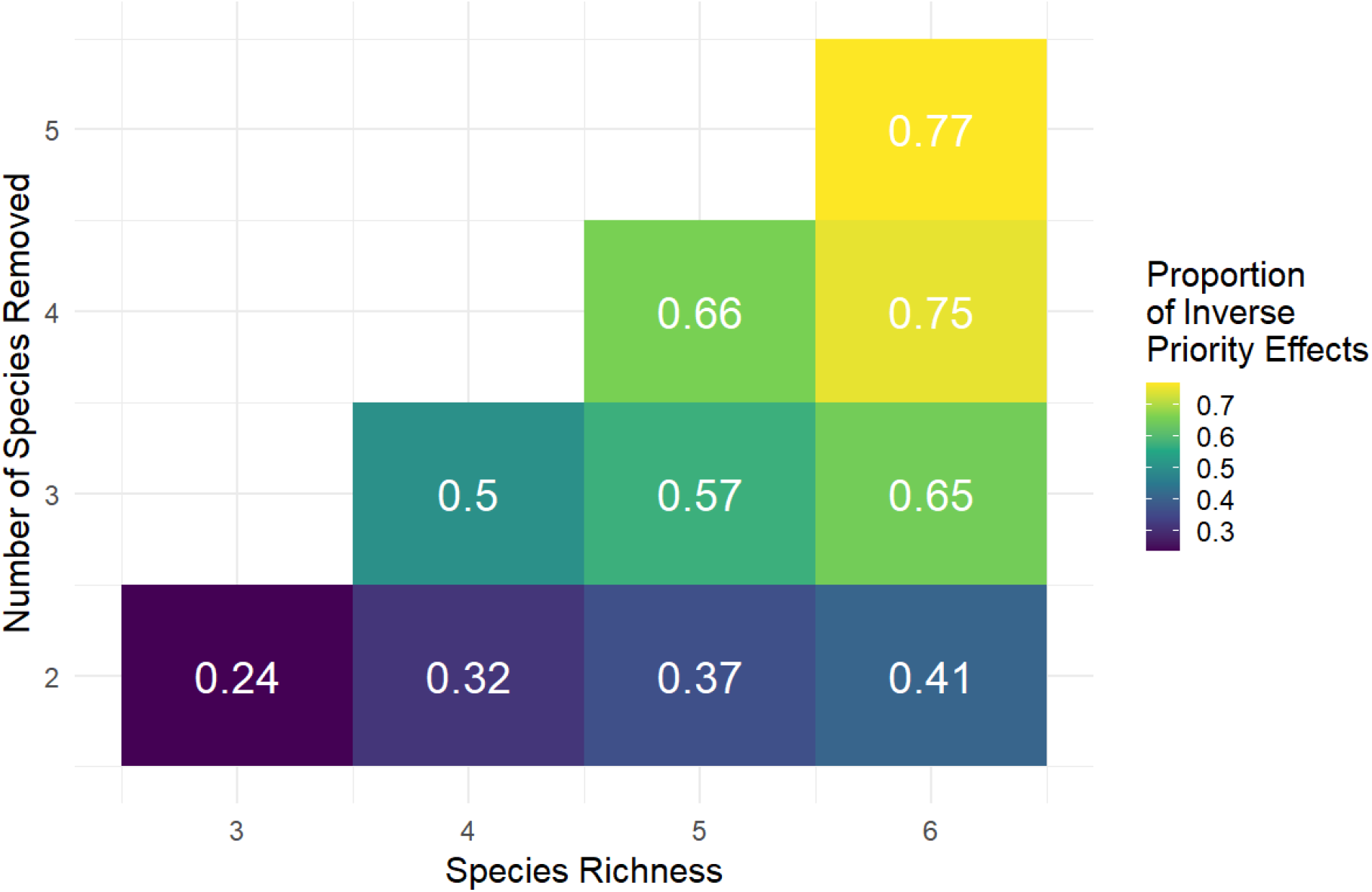
Prevalence of inverse priority effects in competitive communities. Proportion of extinction sequences with inverse priority effect for given species richness and length of extinction sequence in competitive communities.

## H Empirical values for combinations of species richness and length of extinction sequence in communities with competition and facilitation

**Figure H.1:**
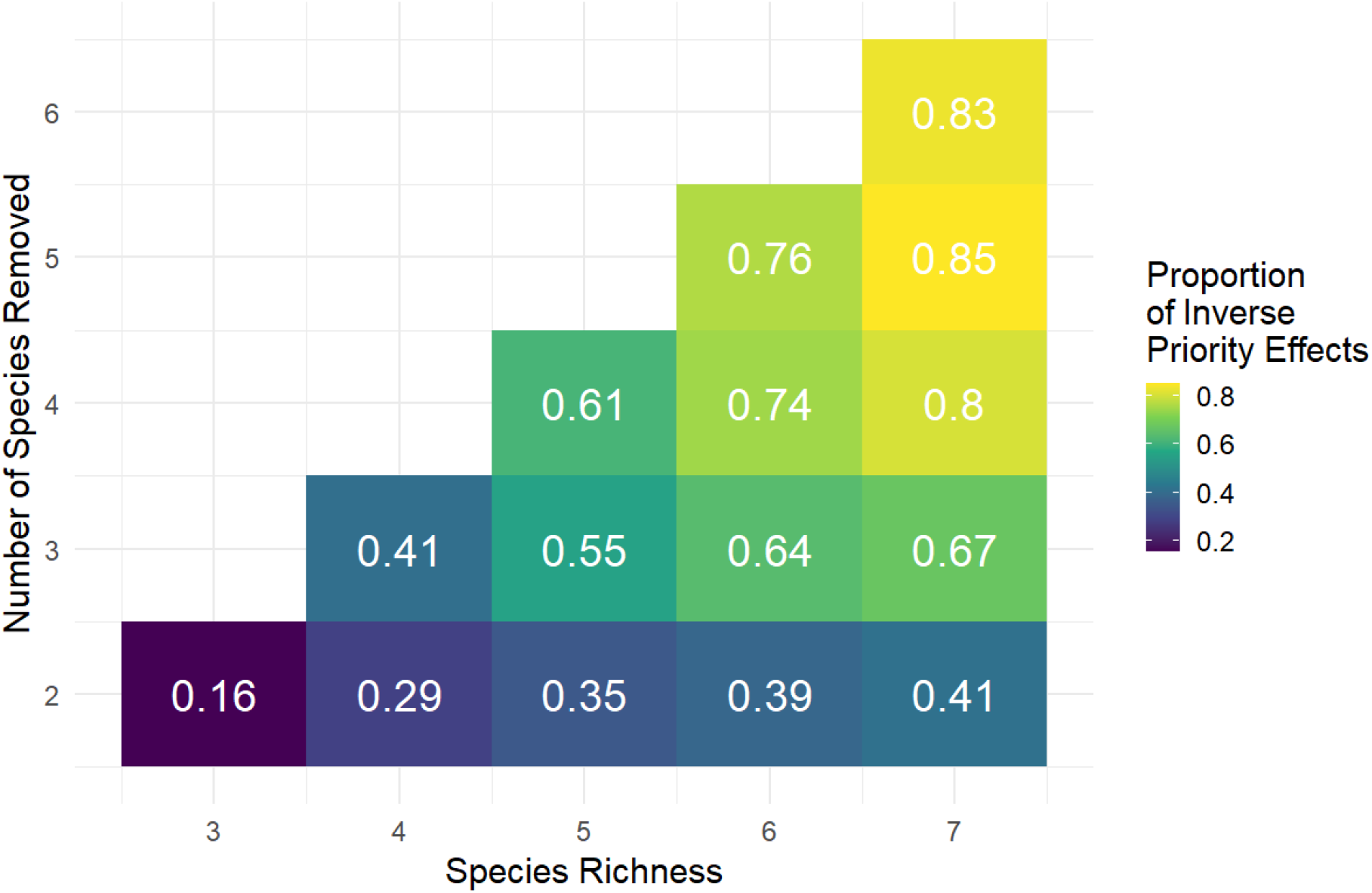
Prevalence of inverse priority effects in communities with competition and facilitation. Proportion of extinction sequences with inverse priority effect for given species richness and length of extinction sequence in communities with competition and facilitation.

## I Locally stable rather than permanent three species communities

If we consider three species competitive communities that coexist with local rather than global stability, we find a change in the relationship between predictors and the presence of inverse priority effects. If we consider only globally attracting three species coexistence states, as asymmetry and mean interaction strength increase, so do the number of extinction sequences that correspond to an inverse priority effect. However, when we require local and not global stability, we gain a new class of dynamics shown in Figure I.1 that does not follow this trend.

**Figure I.1:**
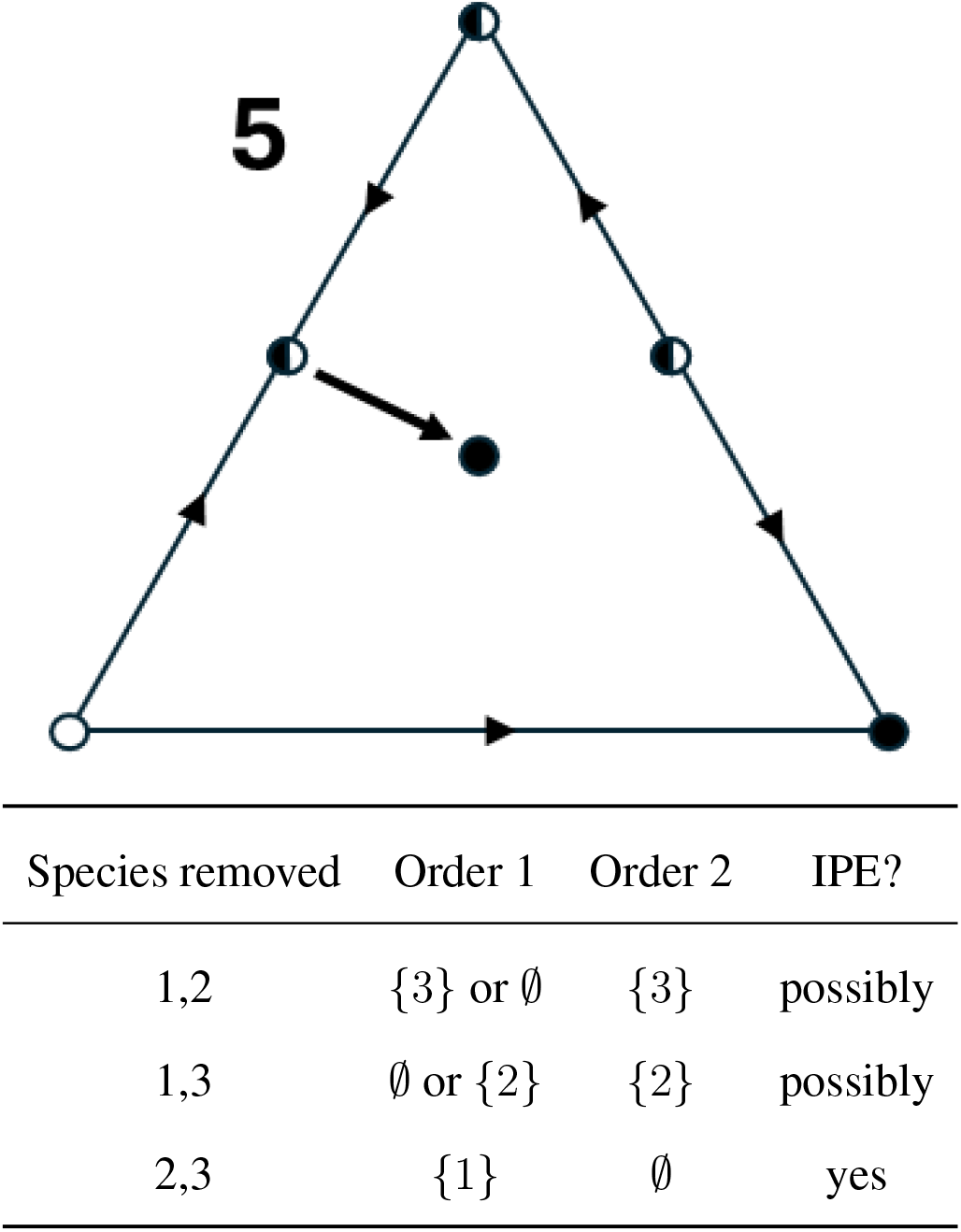
Identifying possible inverse priority effects (IPE) for stable but not permanent three species coexistence state. Bottom left node denotes species 1, bottom right node denotes species 2, and top node denotes species 3. Filled circle denotes an attractor, half filled circle denotes a saddle point, and an unfilled circle denotes a fully unstable equilibrium.

**Figure I.2:**
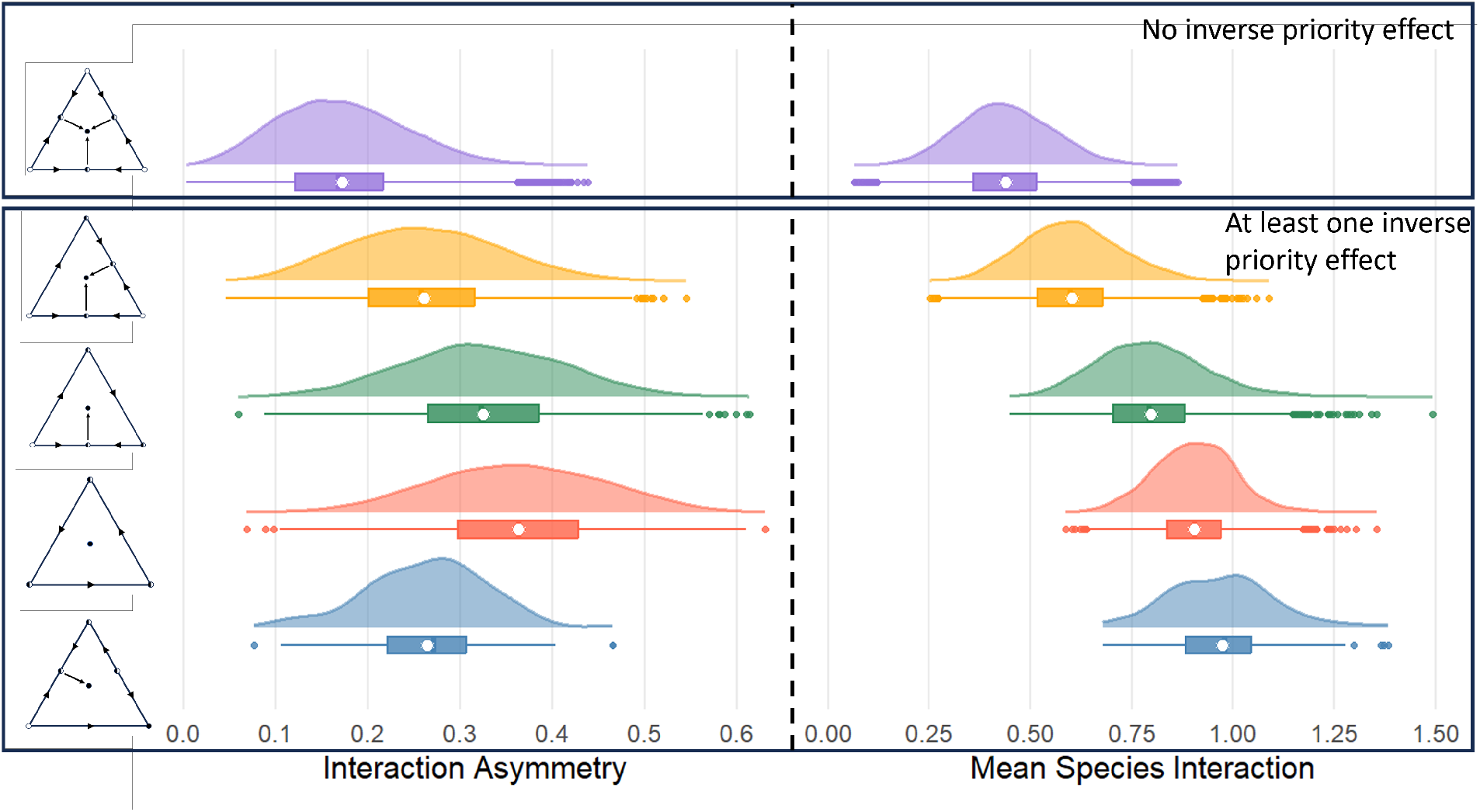
Three species, locally stable communities do not follow the same qualitative trends as permanent communities. Density plots of asymmetry of simulated parameterizations with no inverse priority effects and at least one inverse priority effect. Each row denotes a different dynamic classes 1*−* 5, from top to bottom respectively. The left column shows each class’s interaction asymmetry values while the right column shows the mean species interaction strength. Below each density plot is a boxplot with the white circle denoting the mean value. The last row contains the dynamic class not previously considered.

## References

L. K. Albertson, L. S. Sklar, B. B. Tumolo, W. F. Cross, S. F. Collins, and H. A. Woods. The ghosts of ecosystem engineers: Legacy effects of biogenic modifications. Functional Ecology, 38(1):52–72, 2024.

S. Allesina and S. Tang. Stability criteria for complex ecosystems. Nature, 483(7388):205–208, 2012.

M. I. Ávila-Thieme, E. Kusch, D. Corcoran, S. P. Castillo, F. S. Valdovinos, S. A. Navarrete, and P. A. Marquet. Networkextinction: An r package to simulate extinction propagation and rewiring potential in ecological networks. Methods in Ecology and Evolution, 14(8): 1952–1966, 2023.

G. Barabás, L. Pásztor, G. Meszéna, and A. Ostling. Sensitivity analysis of coexistence in ecological communities: theory and application. Ecology Letters, 17(12):1479–1494, 2014.

S. Berg, A. Pimenov, C. Palmer, M. Emmerson, and T. Jonsson. Ecological communities are vulnerable to realistic extinction sequences. Oikos, 124(4):486–496, 2015.

S. Bewick, B. J. Brosi, and P. R. Armsworth. Predicting the effect of competition on secondary plant extinctions in plant–pollinator networks. Oikos, 122(12):1710–1719, 2013.

C. Borrvall and B. Ebenman. Early onset of secondary extinctions in ecological communities following the loss of top predators. Ecology letters, 9(4):435–442, 2006.

M. C. Bortolan, P. Kalita, J. A. Langa, and R. d. O. Moura. A theoretical and computational study of heteroclinic cycles in lotka–volterra systems. Journal of Mathematical Biology, 90 (3):28, 2025.

J. Brennan and S. J. Schreiber. Using modern coexistence theory to understand community disassembly. Ecology Letters, 29(6):e70424, 2026.

J. F. Brodie, C. E. Aslan, H. S. Rogers, K. H. Redford, J. L. Maron, J. L. Bronstein, and C. R. Groves. Secondary extinctions of biodiversity. Trends in ecology & evolution, 29(12):664–672, 2014.

L. Buche, L. G. Shoemaker, P. Vesk, L. M. Hallett, O. Godoy, and M. Mayfield. Neighbor density-dependent facilitation promotes coexistence and internal oscillation. Ecological Monographs, 95(4):e70040, 2025.

V. Calleja-Solanas, R. O. Moura, J. A. Langa, J. R. Portillo, F. Soler-Toscano, and O. Godoy. A general framework for invasion cycles in ecology. Ecology, 107(4):e70380, 2026.

T. J. Case. Invasion resistance arises in strongly interacting species-rich model competition communities. Proceedings of the National Academy of Sciences, 87(24):9610–9614, 1990.

T. J. Case. Invasion resistance, species build-up and community collapse in metapopulation models with interspecies competition. Biological journal of the Linnean Society, 42(1-2): 239–266, 1991.

P. Chesson. Mechanisms of maintenance of species diversity. Annual review of Ecology and Systematics, 31(1):343–366, 2000.

P. Chesson. Updates on mechanisms of maintenance of species diversity. Journal of ecology, 106(5):1773–1794, 2018.

P. Chesson and J. J. Kuang. The interaction between predation and competition. Nature, 456 (7219):235–238, 2008.

A. T. Clark, L. G. Shoemaker, J.-F. Arnoldi, G. Barabás, R. Germain, O. Godoy, L. Hallett, C. Karakoç, S. Saavedra, and S. J. Schreiber. A practical guide to characterising ecological coexistence. Biological Reviews, 101(1):195–220, 2026.

P. W. Collins, B. C. Latta, and G. W. Roemer. Does the order of invasive species removal matter? the case of the eagle and the pig. PLoS One, 4(9):e7005, 2009.

R. K. Colwell, R. R. Dunn, and N. C. Harris. Coextinction and persistence of dependent species in a changing world. Annual Review of Ecology, Evolution, and Systematics, 43(1): 183–203, 2012.

A. Curtsdotter, A. Binzer, U. Brose, F. de Castro, B. Ebenman, A. Eklöf, J. O. Riede, A. Thierry, and B. C. Rall. Robustness to secondary extinctions: comparing trait-based sequential deletions in static and dynamic food webs. Basic and Applied Ecology, 12(7):571–580, 2011.

I. Donohue, O. L. Petchey, S. Kéfi, A. Génin, A. L. Jackson, Q. Yang, and N. E. O’Connor. Loss of predator species, not intermediate consumers, triggers rapid and dramatic extinction cascades. Global Change Biology, 23(8):2962–2972, 2017.

J. A. Dunne, R. J. Williams, and N. D. Martinez. Network structure and biodiversity loss in food webs: robustness increases with connectance. Ecology letters, 5(4):558–567, 2002.

J. A. Dunne, R. J. Williams, and N. D. Martinez. Network structure and robustness of marine food webs. Marine Ecology Progress Series, 273:291–302, 2004.

C. Emary and D. Evans. Can a complex ecosystem survive the loss of a large fraction of its species? a random matrix theory of secondary extinction. Oikos, 130(9):1512–1522, 2021.

M. S. Fowler. Extinction cascades and the distribution of species interactions. Oikos, 119(5): 864–873, 2010.

M. S. Fowler. The form of direct interspecific competition modifies secondary extinction patterns in multi-trophic food webs. Oikos, 122(12):1730–1738, 2013.

C. M. Godwin, F.-H. Chang, and B. J. Cardinale. An empiricist’s guide to modern coexistence theory for competitive communities. Oikos, 129(8):1109–1127, 2020.

B. Goh. Stability in models of mutualism. The American Naturalist, 113(2):261–275, 1979.

T. N. Grainger, J. M. Levine, and B. Gilbert. The invasion criterion: a common currency for ecological research. Trends in ecology & evolution, 34(10):925–935, 2019.

T. N. Grainger, K. Stark, C. Song, M. A. Barbour, and R. M. Germain. The equilibrium conundrum. Ecology Letters, 28(11):e70232, 2025.

J. Grilli, M. Adorisio, S. Suweis, G. Barabás, J. R. Banavar, S. Allesina, and A. Maritan. Feasibility and coexistence of large ecological communities. Nature communications, 8(1):14389, 2017.

A. Hastings, K. C. Abbott, K. Cuddington, T. Francis, G. Gellner, Y.-C. Lai, A. Morozov, S. Petrovskii, K. Scranton, and M. L. Zeeman. Transient phenomena in ecology. Science, 361(6406):eaat6412, 2018.

J. Hofbauer. Heteroclinic cycles in ecological differential equations. Equadiff 8, pages 105–116, 1994.

J. Hofbauer and S. J. Schreiber. Permanence via invasion graphs: incorporating community assembly into modern coexistence theory. Journal of mathematical biology, 85(5):54, 2022.

J. Hofbauer and K. Sigmund. Evolutionary games and population dynamics. Cambridge university press, 1998.

V. Hutson and R. Law. Permanent coexistence in general models of three interacting species. Journal of Mathematical Biology, 21(3):285–298, 1985.

J. Jiang and F. Liang. Global dynamics of 3d competitive lotka-volterra equations with the identical intrinsic growth rate. Journal of Differential Equations, 268(6):2551–2586, 2020.

R. Kehoe, E. Frago, and D. Sanders. Cascading extinctions as a hidden driver of insect decline. Ecological Entomology, 46(4):743–756, 2021.

T. H. Larsen, N. M. Williams, and C. Kremen. Extinction order and altered community structure rapidly disrupt ecosystem functioning. Ecology letters, 8(5):538–547, 2005.

A. Liaw and M. Wiener. Classification and regression by randomforest. R News, 2(3):18–22, 2002. URL https://CRAN.R-project.org/doc/Rnews/.

R. M. May. Will a large complex system be stable? Nature, 238(5364):413–414, 1972.

R. M. May and W. J. Leonard. Nonlinear aspects of competition between three species. SIAM journal on applied mathematics, 29(2):243–253, 1975.

J. Memmott, N. M. Waser, and M. V. Price. Tolerance of pollination networks to species extinctions. Proceedings of the Royal Society of London. Series B: Biological Sciences, 271 (1557):2605–2611, 2004.

J. Mierczyński and S. J. Schreiber. Kolmogorov vector fields with robustly permanent subsystems. Journal of mathematical analysis and applications, 267(1):329–337, 2002.

E. J. O’Gorman, U. Jacob, T. Jonsson, and M. C. Emmerson. Interaction strength, food web topology and the relative importance of species in food webs. Journal of animal Ecology, 79 (3):682–692, 2010.

F. M. Padilla and F. I. Pugnaire. The role of nurse plants in the restoration of degraded environments. Frontiers in Ecology and the Environment, 4(4):196–202, 2006.

R. T. Paine. Food-web analysis through field measurement of per capita interaction strength. Nature, 355(6355):73–75, 1992.

R. Ranjan, T. Koffel, and C. A. Klausmeier. The three-species problem: Incorporating competitive asymmetry and intransitivity in modern coexistence theory. Ecology Letters, 27(4):e14426, 2024.

P. L. Reynolds, J. Glanz, S. Yang, C. Hann, J. Couture, and E. Grosholz. Ghost of invasion past: legacy effects on community disassembly following eradication of an invasive ecosystem engineer. Ecosphere, 8(3):e01711, 2017.

R. P. Rohr, S. Saavedra, and J. Bascompte. On the structural stability of mutualistic systems. science, 345(6195):1253497, 2014.

G. Q. Romero, T. Gonçalves-Souza, C. Vieira, and J. Koricheva. Ecosystem engineering effects on species diversity across ecosystems: a meta-analysis. Biological Reviews, 90(3):877–890, 2015.

D. Sanders, R. Kehoe, and F. F. van Veen. Experimental evidence for the population-dynamic mechanisms underlying extinction cascades of carnivores. Current Biology, 25(23):3106–3109, 2015.

S. Schreiber, J. Brennan, and J. W. Spaak. From lotka-volterra dynamics to community assembly: Theory, topography, and empirical applications. bioRxiv, pages 2026–07, 2026.

S. J. Schreiber. Criteria for cr robust permanence. Journal of Differential Equations, 162(2): 400–426, 2000.

S. J. Schreiber. Persistence despite perturbations for interacting populations. Journal of Theoretical Biology, 242(4):844–852, 2006.

J. W. Spaak, O. Godoy, and F. De Laender. Mapping species niche and fitness differences for communities with multiple interaction types. Oikos, 130(12):2065–2077, 2021.

G. Strona and C. J. Bradshaw. Co-extinctions annihilate planetary life during extreme environ-mental change. Scientific reports, 8(1):16724, 2018.

T. Therneau, B. Atkinson, B. Ripley, and M. B. Ripley. Package ‘rpart’. Available online: cran. ma. ic. ac. uk/web/packages/rpart/rpart. pdf (accessed on 20 April 2016), 2:5–32, 2015.

A. Torres, S. E. Kuebbing, K. L. Stuble, S. A. Catella, M. A. Núñez, and M. A. Rodriguez-Cabal. Inverse priority effects: A role for historical contingency during species losses. Ecology Letters, 27(1):e14360, 2024a.

A. Torres, T. Morán-López, M. A. Rodriguez-Cabal, and M. A. Núñez. Inverse priority effects: the order and timing of removal of invasive species influence community reassembly. Journal of Applied Ecology, 61(1):51–62, 2024b.

S. Wood and M. S. Wood. Package ‘mgcv’. R package version, 1(29):729, 2015.

M. L. Zeeman. Hopf bifurcations in competitive three-dimensional lotka–volterra systems. Dynamics and stability of systems, 8(3):189–216, 1993.

